# Supervised generative design of regulatory DNA for gene expression control

**DOI:** 10.1101/2021.07.15.452480

**Authors:** Jan Zrimec, Xiaozhi Fu, Azam Sheikh Muhammad, Christos Skrekas, Vykintas Jauniskis, Nora K. Speicher, Christoph S. Börlin, Vilhelm Verendel, Morteza Haghir Chehreghani, Devdatt Dubhashi, Verena Siewers, Florian David, Jens Nielsen, Aleksej Zelezniak

## Abstract

In order to control gene expression, regulatory DNA variants are commonly designed using random synthetic approaches with mutagenesis and screening. This however limits the size of the designed DNA to span merely a part of a single regulatory region, whereas the whole gene regulatory structure including the coding and adjacent non-coding regions is involved in controlling gene expression. Here, we prototype a deep neural network strategy that models whole gene regulatory structures and generates *de novo* functional regulatory DNA with prespecified expression levels. By learning directly from natural genomic data, without the need for large synthetic DNA libraries, our ExpressionGAN can traverse the whole sequence-expression landscape to produce sequence variants with target mRNA levels as well as natural-like properties, including over 30% dissimilarity to any natural sequence. We experimentally demonstrate that this generative strategy is more efficient than a mutational one when using purely natural genomic data, as 57% of the newly-generated highly-expressed sequences surpass the expression levels of natural controls. We foresee this as a lucrative strategy to expand our knowledge of gene expression regulation as well as increase expression control in any desired organism for synthetic biology and metabolic engineering applications.

## 1. Introduction

Gene expression is a fundamental process underlying the cellular functionality of all living organisms and researchers have been trying to control it for decades. A major factor driving our ability to control gene expression arises from our understanding of the cell’s intrinsic regulatory code ^1^, which in turn can be used to design sequences with target expression levels ^2–4^. State of the art machine learning approaches have proven highly useful in this endeavour, helping to expand our knowledge of the DNA regulatory grammar driving gene expression ^5–8^, design novel promoter and gene sequences ^9,10^ and accurately predict gene expression across multiple model organisms ^5,11^. The striking capacity of random DNA to evolve into functioning regulatory sequences by introducing only a couple of bps of mutations, recently shown in bacteria ^12^, suggests that the richness and plasticity of the DNA regulatory grammar results in a vast functional regulatory sequence space far larger than the one existing in nature ^6^. By learning this regulatory sequence space using advanced deep learning approaches ^9,13,14^, one can in principle design systems that precisely traverse it to extract completely novel sequence variants with target expression levels.

Multiple recent studies show that apart from tuning codon usage in gene coding regions ^15,16^, also the DNA sequence of non-coding regulatory regions must be fine-tuned in order to accurately control gene expression ^5,6,17,18^. Proper orchestration of gene expression depends on the interaction of regulatory patterns across the whole *cis*-regulatory structure around the gene, including promoters, terminators, coding and untranslated regions (UTRs) ^1,5^. Despite this, the standard synthetic engineering approach to design regulatory regions of varying expression levels is to apply random mutagenesis in a specific region, most commonly the promoter ^6,19–21^, though also UTRs ^17,22^ and terminators ^23,24^ have been targeted, frequently perturbing only short DNA segments of less than 100 bp. Similarly, knowledge-guided approaches focus on single regions to design minimal synthetic constructs and either stack multiple known highly-functional sequence motifs ^2,4^ or apply machine learning to design them in a generative fashion ^9^. Thus, existing approaches to design DNA sequences have limited control of gene expression, instead relying on experimental screening of large amounts of random synthetic sequences to find functional variants with desired expression levels ^3,6,9^. This inherent ‘blindness’ in relating sequence to expression and the high resource intensiveness, due to the large sequence space that needs to be explored, are also the major factors constraining the length of the explored DNA to only short segments. However, based on recent achievements in modeling DNA and protein spaces ^9,10,14^, we hypothesize that state-of-the-art generative deep neural networks are capable of learning the entire DNA regulatory landscape directly from natural genomic sequences. Coupled with leveraging information from the whole gene regulatory structure including the coding region ^1,5^, this can not only help to overcome the existing experimental limitations, but can also enable precise and gene-specific navigation of the regulatory sequence space, boosting the accuracy of expression control by generating *de novo* regulatory DNA with desired expression levels.

In the present study, we use deep learning frameworks to demonstrate that a generative modeling approach can successfully design novel yet functional regulatory DNA in *Saccharomyces cerevisiae*, outperforming targeted mutational approaches in mRNA expression optimization. The deep neural networks are trained only on natural genomic sequences spanning the whole gene regulatory structure comprising the promoter, UTRs and terminator. First, we verify that a conventional mutagenesis approach with *in silico* screening, (i) using a highly accurate deep predictive model ^5^ and (ii) including targeted mutagenesis of only the most relevant DNA positions, is inefficient at generating novel functional sequences. Next, we apply deep generative adversarial networks to design *de novo* gene regulatory sequences with natural-like properties. Using an optimization procedure that couples the generative and predictive neural networks ^5,14^, we add coding sequence information to the generative approach and learn to precisely navigate the regulatory sequence-expression landscape of a specific gene across almost 6 orders of magnitude of expression levels, accurately controlling the sampling of sequences with targeted expression levels. Sequence properties of the generated regulatory DNA, including *cis*-regulatory grammar such as DNA motifs and motif associations, reflect those of natural sequences across the range of expression levels. In fact, the generated sequences retain a natural or even higher level of dissimilarity (>30%) to any currently known regulatory sequence. Finally, we experimentally verify the generated constructs and find that experimentally measured mRNA expression levels reflect predicted ones across 3 orders of magnitude, with 57% of the constructs designed to be highly expressed surpassing the level of gene expression of natural high-expression control sequences.

## 2. Results

### Random mutagenesis requires multiple testing rounds

Driven by the idea that DNA sequences are predictive of gene expression levels ^1,5,11^, we reasoned that randomly mutating DNA sequences coupled to virtual screening would be a plausible strategy for gene expression control. To design sequences with increased or decreased gene expression levels, we first set up a random mutagenesis approach with *in silico* screening (Methods M3) using an experimentally validated highly-accurate predictive model (predictor, *R^2^_test_* = 0.8) of yeast gene expression ^5^ trained on natural genomic sequences comprising whole gene regulatory structures of 1000 bps (Figure 1A,B, Figure S1, Methods M1,2). We focused on mutating the promoter region spanning 400 bp (previously found as the optimal predictive region size ^5^), whereas the other regions (UTRs and terminator) were kept fixed. Apart from the initial strategy of blindly mutating whole 400 bp promoter regions, as an additional strategy, we used the predictor to inform the mutational procedure by querying its sensitivity to specific positions in the promoter sequence (Figure 1B, Methods M3). Here, only the most sensitive and thus relevant positions were preferentially used as the scaffolds for targeted mutagenesis (77 bp on average, Figure S2).

**Figure 1.**
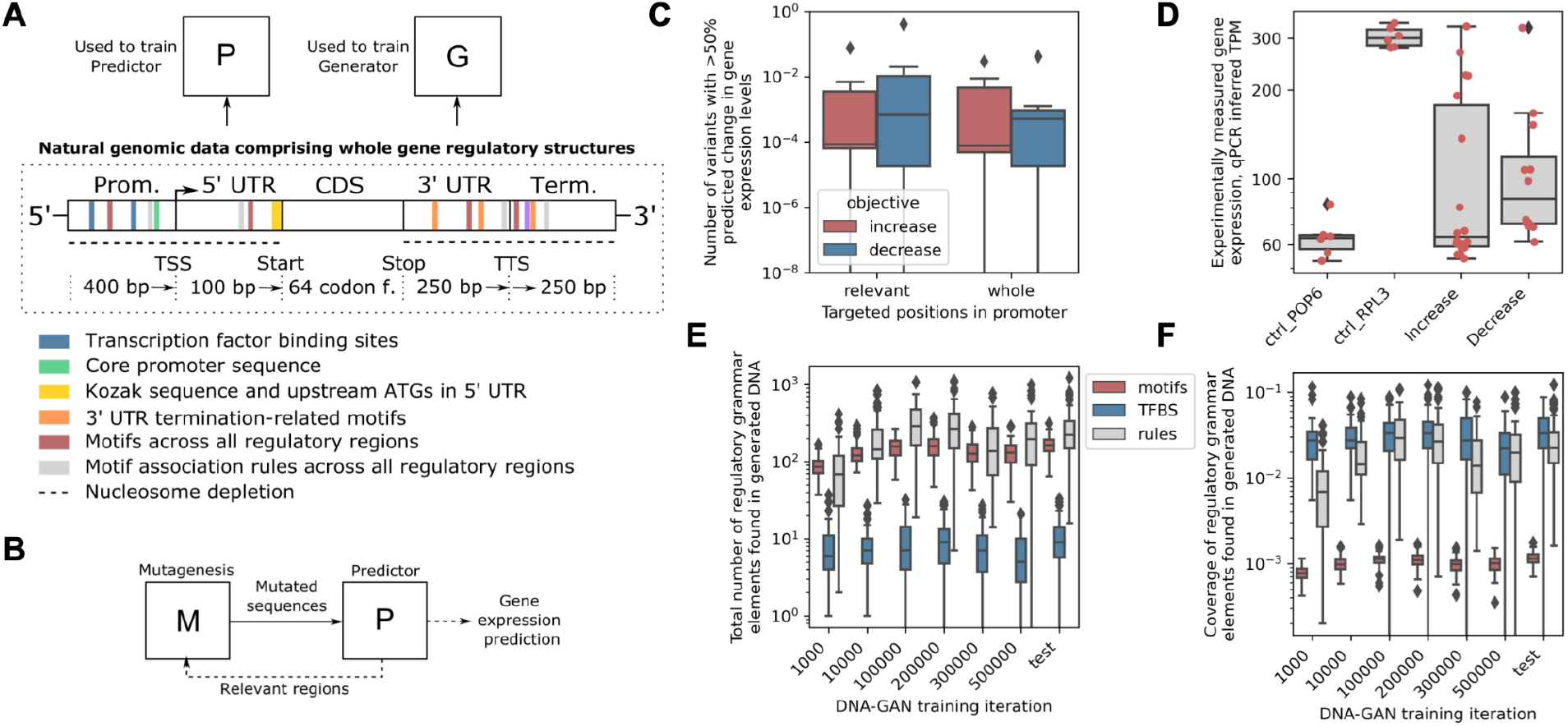
Implementing mutational and generative strategies to design regulatory DNA. (A) Schematic depiction of the *Saccharomyces cerevisiae* natural genomic sequencing dataset that was used to train both the predictive (P) ^5^ and generative (G) models used in the study. The dataset spanned the whole gene regulatory structure of 1000 bps and included promoter, terminator and untranslated regions (UTRs) as well as codon frequencies of coding regions. Marked are the different natural sequence properties related to DNA *cis*-regulatory grammar that were further analysed with the generator. (B) Schematic depiction of the mutagenesis strategy that included *in silico* screening, where a random mutagenesis procedure (M) was coupled with a predictor (P) of yeast gene expression ^5^, which was also used to inform the mutational procedure on which positions were the most relevant to mutate (Methods M3). (C) Amount of mutated sequence variants that achieved an over 50% increase or decrease in predicted gene expression levels by mutating 10% (40 bp) of whole promoter regions (400 bp) or only the most relevant promoter positions. (D) Quantitative PCR (qPCR) measurements of mRNA levels with 10 mutated RPL3 sequence variants predicted to achieve ~2-fold increases or decreases in expression levels from the native regulatory sequence (see Table S2). Apart from the native regulatory regions of RPL3 (predicted TPM of 303), POP6 regions were used as a low control (predicted TPM of 64). (E) Total number and (F) coverage of transcription factor binding sites (TFBS), DNA motifs and motif association rules ^5^ uncovered in samples of generated or natural test sequences across generator training iterations.

We evaluated the mutagenesis approach by creating and assessing 16.8 million sequence variants at different parameters using 7 natural regulatory regions as scaffolds (Figure S3, Methods M3). When aiming to achieve an over 50% increase or decrease in mRNA expression levels, we found that on average, at most 0.3% of the sequence variants were predicted to achieve the desired effect when mutating 10% (40 bp) of whole promoter regions, which increased to 0.4% when mutating the most relevant promoter regions (Figure 1C). Unsurprisingly, this value greatly decreased with lower percentages of mutated sequence size (Table S1). We then selected and experimentally tested 10 of the mutated regulatory sequence variants of the RPL3 gene with the largest predicted (~2-fold) increase or decrease from the native levels, including both whole and only relevant-region mutational strategies and different percentages of mutated sequence size (5 and 10%, Table S2, Methods M5,6). Of the tested variants, 40% corresponded with predictions, of which all were designed to decrease expression (Figure 1D, Figure S4). This indicates that even fewer sequences than the above computational estimations are functional when designed by the random mutagenesis approach, thus still requiring multiple rounds of selection and testing despite the use of *in silico* screening.

### Deep generative modeling of regulatory DNA

The above results suggested that, in order to have a more controlled approach of designing synthetic regulatory DNA, alternative strategies to random mutagenesis are required and should be explored. We therefore tested if an altogether different, generative modeling strategy could be used to design regulatory sequence variants with increased or decreased expression levels, by learning the genetic regulatory and expression landscape directly from natural genomes. We trained a generative model (generator) using whole gene regulatory structures (Figure 1A) with a generative adversarial network (GAN) approach ^25^, where a discriminator network was used to train a generator, both comprising 6 convolutional layers (Figure S5, Methods M4). As input data from which to learn the distribution of the gene regulatory sequence space, we used 4238 sequences of whole gene regulatory structures from yeast, previously found to span all the regulatory features important for predicting over 82% of the variability of mRNA expression levels ^5^ (Methods M1). The performance of the generator was computationally validated by verifying that the sequence properties of the generated variants reflected those of natural sequences, including: (i) sequence compositional validity, (ii) sequence similarity measures, (iii) predicted gene expression levels and (iv) known *cis*-regulatory grammar, per generated sequence (Figure 1A, Table S3, Methods M7). Indeed, after training, the majority of the generated sequences (86%) displayed natural sequence-like properties (Figure 1E,F, Figure S6), containing not only appropriate sequence composition but also known DNA regulatory motifs ^1^ including Jaspar ^26^ and Yeastract ^27^ transcription factor binding sites (TFBS), and motif associations ^5^ (Figure 1A). We also verified that the generated sequences retained a sequence diversity similar to that of natural sequences, with the sequence identity of both the generated and test datasets to the train dataset equalling ~67% (Figure S6) and showing that the nucleotide composition of generated variants was as variable and dissimilar to natural sequences as they are amongst themselves (Figure S7). This ensures that the model did not overfit to the training dataset and shows that it can generate *de novo* regulatory sequences with properties indistinguishable from natural ones across a wide range of expression levels.

### Precise gene-specific navigation of DNA regulatory sequence-expression landscape

Next, in order to explore the generative model in a directed-evolution fashion and devise a procedure that produces regulatory sequences with target expression levels, we set up an optimization procedure ^14^ (Figure 2A). The trained DNA regulatory sequence generator and gene expression-predictor ^5^ neural networks were coupled in a feedback loop, where the predictor was used to optimize the expression level of generated sequences by guiding the latent space of the generator, thus defining the input variables used to draw new sequence samples (Figure 2B, Methods M4). Since the predictor also evaluates variables describing the coding region (see Figure 1A: 64 coding frequencies), this procedure in fact couples the generated regulatory structures to a specific gene of interest. Merging the results of both maximization and minimization of gene expression and using t-distributed stochastic neighbour embedding (t-SNE) dimensionality reduction ^28^ over the latent vectors confirmed that with this approach, desired expression levels are mapped to identified latent subspace resembling a continuous multidimensional curve that covers ~6 orders of magnitude of expression levels (Figure 2C, Figure S8). Thus, with optimization, the dynamic range of expression levels of the generated sequences increased over 3-fold compared to those obtained by randomly sampling the generator (in equally sized samples), even surpassing the natural range of expression levels for a specific gene of interest (Figure 2C: GFP coding sequence shown). Similarly as before, analysis of sequence identity verified that the sequences produced by the generator optimization were not similar to any natural ones and retained the natural sequence diversity of ~0.67 (Figure S9, Methods M7). Importantly, by sampling and computationally analysing sequence selections across a 4 order-of-magnitude range of expression levels (see Figure 1C, Methods M4), generated variants displayed sequence properties (Figure 1B, Table S3, Methods M7) reflecting those of natural sequences and indicating that they are potentially functional (Figure 2D,E, Figure S10). For instance, the overall amounts of *cis*-regulatory grammar were observed to steadily increase in proportion to the predicted gene expression levels (Figure 2D,E). This suggests that points in the generator’s latent space with desired expression levels can be sampled, which generalize beyond the naturally available regulatory sequence space to generate novel but functional sequence diversity.

**Figure 2.**
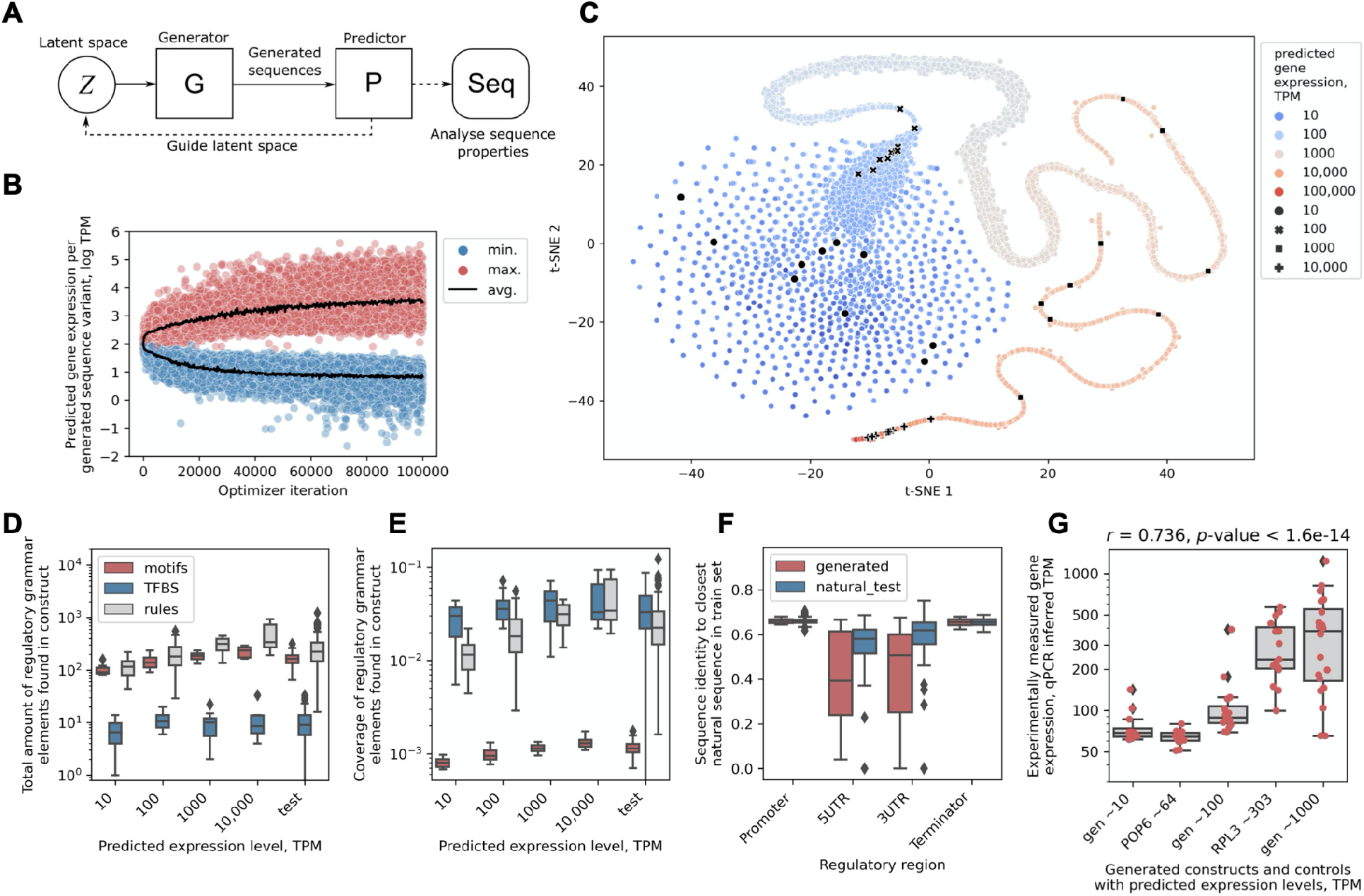
Predictor-guided generator optimization enables precise and gene-specific navigation of the regulatory sequence-expression landscape. (A) Schematic depiction of the procedure to optimize the generator using a trained predictor ^5^, which introduces coding region information into the generative approach and explores the input latent space of the generator to produce sequence variants across the whole gene expression range, providing precise navigation across the gene regulatory sequence-expression landscape. (B) Predicted expression levels of generated sequence variants across optimization iterations set to either maximize or minimize expression levels. (C) T-distributed stochastic neighbour embedding (t-SNE) ^28^ mapping of the input latent subspaces that produce novel sequence variants spanning ~6 orders of magnitude of gene expression (colored and black dots), uncovered using the predictor-guided generator optimization. Black dots represent selections of 10 sequence variants per each of 4 expression groups covering a 4 order-of-magnitude range of predicted expression levels from TPM ~10 to ~10,000. (D) Total number and (E) coverage of transcription factor binding sites (TFBS), DNA motifs and motif association rules ^5^ in the groups of sequence variants produced by generator optimization across 4 orders of magnitude of predicted expression levels and natural test sequences. (F) Sequence homology of the generated and test dataset sequences to the respective closest representative sequences in the training dataset across the 4 regions of the gene regulatory structure. (G) Quantitative PCR (qPCR) measurements of mRNA levels with groups of generated sequence variants across 3 orders of magnitude of predicted expression levels (TPM of ~10, ~100 and ~1000, see Table S4). Natural regulatory regions of the POP6 and RPL3 genes were used as low and high controls with a predicted TPM of 64 and 303, respectively.

### *In vivo* gene expression control using generated regulatory DNA

Finally, in order to test the efficacy and validity of the generative procedure, we selected a group of generated sequences across a 3 order-of-magnitude range of predicted expression levels (Figure 2C: TPM of ~10, ~100 and ~1000). This was based on a selection procedure that retained sequences with properties that corresponded to those of natural sequences in each respective expression range (Figure 2D,E, Figure S10, Methods M4), with the exception that sequences with the lowest sequence similarity to natural ones were preferred (i.e. most different from natural, Figure 2F, Figure S11. Similarly, in each expression range, sequence diversity was maximised (meaning no 2 tested sequences were alike, Figure S11), in order to test a wide range of unique sequence variants and not merely multiple versions of a common variant. Although mRNA levels were measured, the GFP gene was used due to its low effect on cell growth (Figure S12). As a result of the limitations imposed by sequence synthesis, limiting the possibility to synthesize and test very highly or lowly expressed sequences (e.g. <10 and >1000 TPM), we succeeded in testing 17 regulatory sequence-GFP constructs. Additionally, regulatory structures of the well known POP6 (predicted TPM of 64) and RPL3 (predicted TPM of 303) genes ^5^ were used as low and high controls, respectively.

We observed that experimental measurements of the mRNA levels produced by each construct achieved strong correlation with the predicted levels (Pearson’s *r* = 0.74, *p*-value < 1.6e-14, Figure S13, Methods M6). The measured levels were also significantly correlated with specific sequence properties related to DNA regulatory grammar, including the number and coverage of motifs and motif association rules ^1,5^, 5’ UTR length ^7,29^, nucleosome depletion ^30,31^ as well as strong Kozak ^32,33^ and positioning elements ^4,34^ (Pearson’s *r* > 0.38, *p*-value < 7.6e-3, Figure S14). Despite the strong correlation, on average, the measured expression levels reflected the predicted expression range only in the group of constructs with the predicted TPM of ~100 (Figure 2G: avg. measured TPM of 114), whereas a 7.7-fold and 2.5-fold difference between predictions and measurements was observed with the lower (predicted TPM ~10, avg. measured TPM 77) and higher groups (predicted TPM ~1000, avg. measured TPM 397), respectively. Nevertheless, although we were not able to generate sequences with expression lower than the POP6 control, within the highest expression group, 4 out of 7 regulatory constructs (57%) displayed average expression levels that surpassed those of the natural RPL3 control by up to 2.7-fold (Figure 2F, Table S4). This demonstrates that our generative procedure can independently design completely *de novo* gene regulatory sequences that exceed natural highly expressed controls by learning the natural regulatory DNA variation directly from genomic data, showcasing its usefulness to expand the available supply of synthetic regulatory DNA with novel and improved sequence variants.

## 3. Discussion

In the present study, we asked whether whole *de novo* functional DNA regulatory structures producing target gene expression levels can be generated just from the knowledge of natural regulatory sequences. As there are >10^60^ ways to construct a mere 100 bp promoter sequence, many of which can be functional ^12^ and spanning more DNA variation than exists in all living species on our planet, experimentally exploring even a tiny fraction of such an enormous sequence space is challenging and often infeasible due to the vast species diversity and complexity of eukaryotic gene regulation. Here, we thus used state-of-the-art deep learning algorithms ^5,14,25,35,36^ to learn and map the functional DNA regulatory sequence space to gene expression levels directly from natural genomic data in *Saccharomyces cerevisiae*, enabling the design of novel expression systems in a controlled manner.

This is made possible by multiple development steps and improvements that have enabled us to develop a generative modeling approach, which include: (i) whole gene regulatory structures ^1^ within optimized natural genomic datasets ^5^ (Figure 1A), (ii) highly-accurate predictive models of gene expression levels that can explain over 82% of their variation from regulatory sequence alone ^5^, (iii) deep generative modelling procedures that are capable of learning and expanding functional coding ^10^ and regulatory ^9,14^ sequence spaces from natural genomic data (Figure S5), and (iv) optimization procedures ^14,35,36^ that allow us to include coding region information in sequence design and thus enable gene-specific fine-tuning of generators across the whole range of expression levels (Figure 2A,C). With the latter, due the possibility to connect deep neural networks in end-to-end differentiable architectures, the initial capability of deep generative models to learn the DNA regulatory sequence space from natural genomic data ^9^ is expanded using optimization guided by predictive models (Figure 2A). This enables us to gain control over the generator’s mapping of the regulatory sequence space to the respective gene expression levels and navigate the functional regulatory sequence-expression landscape, to produce generated sequences with desired expression levels for any given gene (Figure 2B,C). We can thus design completely ‘alien’ variants of gene regulatory structures (Figure 2F) that are nevertheless functional, containing natural-like properties and *cis*-regulatory grammar (Figure 2D,E) and even surpassing the expression level of natural highly-expressed genes (Figure 2G). Moreover, since our DNA-generator learns the whole functional regulatory sequence space, it can generate a practically infinite supply of new sequence samples, while traversing only the most relevant subspace instead of sampling random sequences from all 4^1000^ possible variants that would otherwise be needed to explore the 1000 bp of regulatory DNA. Therefore, by advancing generative models to span complete gene regulatory structures and by mapping sequence production directly to the whole range of expression levels, we create the most inclusive solution for gene expression control to date.

We experimentally demonstrate that the generative strategy to design regulatory DNA is more efficient than a mutational one when using models trained purely on natural genomic data. Random mutagenesis is a brute-force strategy that starts with an existing natural scaffold and can only traverse the sequence-expression landscape non-intelligently, a set of mutations at a time. It requires multiple experimental trials to develop functional sequences, even when coupled with predictive models that can accurately map regulatory sequence to gene expression ^5^ (Figure 1B,C,D). The approach is ‘blind’ in its capacity to change sequence content, except for the computational and experimental screening processes that test the functionality of the designed sequences, but which are decoupled from the sequence design. The mutagenesis approach could thus possibly be enhanced using evolutionary optimization algorithms ^37,38^ that would guide the targeted sequence evolution indirectly, requiring the optimization algorithm to explore the predictor’s coupling between generated sequences and expression levels via numerous iterations. In essence, this would be similar to the generator optimization approach used here, however the latter approach presents multiple improvements. Specifically, the generator models the whole functional regulatory landscape at once, producing natural-like functional sequence variants and not merely randomly mutated variants of existing sequences. It is therefore based on two knowledge-based models (Figure 2A: both generator and predictor) and not merely a single one (Figure 1B: only predictor). Due to this, the generative approach explores the whole allowed sequence space and is not limited to exploring mere subspaces that contain also invalid sequence variants, such as when using random mutagenesis. Apart from optimization, informing the mutational procedure by constraining the mutated positions to only relevant ones (Figure 1B), such as those specified by the predictor that contain important binding sites ^5^, might also be an insufficient strategy to improve mutagenesis. This is due to the large number of position-specific interactions for each single nucleotide position that affect protein-binding ^21,39^, which are spread beyond only the most important binding sites and their immediate vicinity ^1,6,9^. For instance, constraining mutagenesis to the surrounding bases of the −35 and −10 promoter binding sites in *E. coli* led to producing very little functional variants ^9,40^, suggesting that the sequence beyond these regions contains important information for generating functional promoters. Similarly, we observed only a very small increase in the capacity to create sequences with increased or decreased expression levels when mutating only relevant positions (Figure 1C). On the other hand, generative models resolve the problem of relevance by mapping positional interactions across whole sequences, learning which positions are the most important for binding and functionality ^9,10^.

In synthetic biology, alternative bio-manufacturing hosts offer several potential benefits for speeding up bioprocess development ^41^, bringing new drug candidates to the clinic and maximizing the use of manufacturing facilities during a pandemic ^42^ or facilitating high-yield manufacturing^43^. To reach desired expression levels that are predictable, robust, and tuneable, genetic constructs that can only be built from well-characterized gene regulatory parts are required ^44^. Moreover, when expressing a gene of interest, all regulatory regions have been shown to affect gene expression levels. For instance, the promoter can be strongly dependent on the choice of the terminator ^5,45^, and both are gene-context dependent and have to be matched with the coding region comprising codon usage that is optimal to facilitate expression ^15,46,47^. While there are tens of thousands of sequenced genomes, our capability to develop such alternative expression systems is highly underdeveloped^48^ and primarily limited by the costly experimental screening approaches used to design and characterize short parts of single genomic regions ^2,6,9^ which is remain challenging for many industrially important strains with low transformation efficiency^49^. Considering the benefits of generative models as well as the costs and resource requirements of synthetic library construction and testing, the use and further development of mutagenesis for regulatory sequence design may not be worthwhile, apart from exploring the intrinsic functionality of expression regulation ^6^. Instead, we demonstrate that the generative approach can produce whole gene regulatory structures, while taking into account also the information from the coding sequence, thus mimicking whole natural regulatory systems in order to ensure the highest control over gene expression. The advantages of the proposed approach are that (i) it requires only natural genomic data as input, with no need for library construction and costly experimental screening, (ii) it can be expanded to virtually any sequenced organism including those with low transformation efficiency, and (iii) it can be used to produce condition-independent or, if required, even condition-dependent regulatory sequences with controllable gene expression that exceeds the expression levels of natural DNA. Therefore, we foresee this as a highly versatile and lucrative strategy to expand our knowledge of gene expression regulation as well as increase expression control in synthetic biology and metabolic engineering applications.

## 4. Methods

### M1. Data

*S. cerevisiae* S288C genome sequence data, including gene sequences, as well as transcript and open reading frame (ORF) boundaries, were obtained from the Saccharomyces Genome Database (https://www.yeastgenome.org/) ^50,51^ and additional published transcript and ORF boundaries were used ^52,53^. Coding and regulatory regions were extracted based on the transcript and ORF boundaries. DNA sequences were one-hot encoded, untranslated region (UTR) sequences were zero-padded up to the specified lengths (Figure 1A: promoter of 400 bp, 5’ UTR of 100bp, 3’ UTR of 250 bp and terminator of 250 bp) ^5^ and the 64 codon frequencies were normalized to probabilities.

For gene expression levels, processed raw RNA sequencing Star counts were obtained from the Digital Expression Explorer V2 database (http://dee2.io/index.html) ^54^ and filtered for experiments that passed quality control, yielding 3025 high-quality RNA-Seq experiments. Raw mRNA data were transformed to transcripts per million (TPM) counts ^55^ and genes with zero mRNA output (TPM < 5) were removed. Prior to modeling, the mRNA counts were Box-Cox transformed ^56^ with lambda set to 0.22. As the mRNA counts and ORF lengths were significantly correlated due to the technical normalization bias from fragment-based transcript abundance estimation ^57^, we computed mRNA counts uncorrelated to gene length. For this, the residual of a linear model, based on ORF lengths as the response variable and mRNA counts as the explanatory variable, was used as the corrected response variable ^5^.

To obtain training datasets, we considered that for the initial 4,975 protein-coding genes with genomic sequence information, median expression levels across the RNA-Seq experiments varied within 1 relative standard deviation (*RSD* = *σ*/*μ*) for 85% of the genes ^5^. We therefore used DNA sequences of the regulatory and coding regions of these 4,238 genes with RSD <1 for training. For predictive modeling, the data comprised paired gene regulatory structure explanatory variables and mRNA count response variables, where a total of 3,433 gene data instances were used for training the model, 381 for tuning the model hyperparameters and 424 for testing. For generative modeling, a total of 3,814 regulatory structure sequences were used for training and the remaining 424 were used as unseen test data. Here, the data was balanced prior to training by distributing the corresponding mRNA counts across 30 bins and sampling input sequence data from all bins such that all the values were uniformly represented instead of using the initial distribution (Figure S15: approximately normal for the Box-Cox transformed data shown).

### M2. Deep predictive modeling

To train a predictive model that predicts gene expression levels from whole gene regulatory structure data, a deep neural network architecture of 3 CNN layers and 2 dense (FC) layers was used ^1,5,58,59^. The network was trained consecutively, first on regulatory sequences input to the first CNN layer and then the dense layers were replaced and the whole network retrained using the numeric variables (codon frequencies) appended to the output of the last CNN layer and input to the first dense layer. Batch normalization ^60^ and weight dropout ^61^ were applied after all layers and max-pooling ^62^ after CNN layers. The Adam optimizer ^63^ with mean squared error (MSE) loss function and ReLU activation function ^64^ with uniform ^65^ weight initialization were used. In total, 26 hyper-parameters were optimized using a tree-structured Parzen estimators approach via Hyperopt v0.1.1 ^66^ at default settings for 1500 iterations with the same initial value ranges as in ^5^. The best models were chosen according to the minimal MSE on the validation set with the least spread between training and validation sets. The coefficient of determination (*R*^*2*^) was defined as *R*^*2*^ = 1 − *SS*_*Residual*_/*SS*_*Total*_[Eq. 1], where *SS*_*Residual*_ is the sum of residual squares of predictions and *SS*_*Total*_ is the total sum of squares, and statistical significance was evaluated using the two-tailed *F*-test. For building and training models Keras v2.2 and Tensorflow v1.10 software packages were used and accessed using the python interface.

### M3. Mutagenesis approach

To design novel regulatory sequence variants with the mutation procedure, either whole promoter sequences or only the most relevant positions were randomly mutated at different settings for the percentage of the mutated sequence size: 1% (4 bp), 2% (8 bp), 5% (20 bp) and 10% (40 bp). This was done while verifying that all mutated variants were different from any of the natural sequences, thus the mutation size also corresponded to the distance from the closest natural sequence. The maximum mutation size of 10% was used in order to limit using the predictor too far outside of its operational range, defined by the natural training sequence space, which can potentially cause incorrect predictions ^67,68^. 300,000 mutations were performed per each of the eight settings per gene scaffold sequence.

To obtain the most relevant positions in the promoter sequence, we calculated the relevance profiles that give an estimate of the sensitivity of the predictive model at specific positions in the input sequence. To calculate the relevance of the different DNA positions for model predictions, defined as *Relevance* =(*Y* − *Y* _*occluded*_) /*Y* [Eq. 1], where *Y* is the model prediction, an input dataset with sliding window occlusions was used with the predictive model to obtain predictions ^69,70^ (Figure S16). The window size of the occlusions was set to either 1 or 10 bp. To obtain only highly sensitive regions, relevance z-scores above a cutoff of 1 were selected.

To calculate the amount of mutated sequence variants that achieved an over 50% increase or decrease in predicted gene expression level, regulatory sequence scaffolds from the following 7 genes were used: YDR541C, POP6, PMU1, YBL036C, MNN9, RPC40, RPL3. Experimental sequence selection was performed with the RPL3 gene, where the mutated sequences were sorted and selected based on largest achieved increases and decreases, targeting ~2-fold changes, as well as according to the limitations imposed by DNA sequence manufacturers. Thus, only sequences with a mutated sequence size of 5 and 10% were selected, where either the whole promoter or only relevant positions with a window size of 10 bp were mutated. When selecting for increased expression while mutating whole promoters, only sequences with a mutated size of 10% achieved the targeted changes in predicted expression levels. Two representatives were selected for increased gene expression per combination of settings and a single representative for decreased expression, yielding the 10 tested sequence constructs (Table S2).

### M4. Deep generative modeling

To devise a system to generate realistic DNA regulatory sequences corresponding to the whole gene regulatory structure, we trained a generative model using a generative adversarial network (GAN) approach ^25^ (Figure S6). In order to capture all the levels of regulatory information across the input sequences, both the generator and discriminator equally comprised 6 convolutional neural network (CNN) layers of opposite orientation, where the first (last) 5 layers were residual blocks containing skip connections with a residual factor of 0.3 ^14,71^. Each CNN layer comprised 100 filters, a kernel size of 5 and a stride of 1. The dense layer size was equal to the input sequence size (1000) x CNN filter size (100). The Adam optimizer ^63^ with the Wasserstein loss function (WGAN) ^72,73^ and ReLU activation function ^64^ with uniform ^65^ weight initialization were used. The learning rate parameter was set to 1e-5, beta1 to 0.5 and beta2 to 0.9, and the batch size was 64. The ratio of discriminator to generator updates was set to 5. The dimensionality of the latent space was set to 200 after testing GANs with 100, 200 and 1000 dimensional latent spaces and finding no improvement in performance over this size, showing that it sufficiently captured the key information in the DNA sequence data. The latent space was sampled according to a standard normal distribution during training.

To generate sequences that manifest desired target expression levels by connecting the functional regulatory DNA space modeled by the generator with expression levels and coding sequence information modeled by the predictor, a DNA-based activation maximization approach ^14,35,36^ was used that incorporates both the trained generator and predictor models (Figure 2A). The optimal trained generative model to use for optimization was identified at iteration 200,000 (Figure S6), further supported by comparing the properties of generators obtained at 6 different training iteration checkpoints (100,000, 200,000, 300,000, 500,000, 700,000 and 1,000,000) after optimization, which included the range of predicted gene expression levels and percentage of unique generated sequences (Figure S17). Optimizations were run for 100,000 iterations and, to increase the breadth of the investigated latent subspace, 10 optimization runs were performed with different initial random states. The results were merged to obtain a set of 6,062,804 unique sequences that were used for further analysis.

To obtain a selection of sequences for experimental validation, the following selection procedure was used. Four expression bins were defined to cover a 4 order of magnitude range of expression levels within a 10% range above or below the TPM values of 10, 100, 1000 and 10,000. Approximately 100 sequences per expression bin and per optimization seed were randomly selected from the above merged optimized sequence dataset, yielding 5706 sequences. Next, by comparing 16 sequence properties (see underlined properties in Table S3) of the generated sequence selection to those of natural test sequences, 452 sequences were subselected with all tested sequence properties within the ranges defined by natural test sequences. From here, the experimental set was constructed by randomly selecting 10 sequences in each expression bin, by optimizing for the highest sequence diversity within each expression bin, whilst retaining the natural sequence diversity (Figure S10: avg. seq. id. of 0.67). The final set of 40 selected sequences was thus highly diverse and as different from natural sequences as these are among themselves, representing as yet unseen sequence variants. Further limitations with sequence synthesis when ordering the selected generated variants as gene fragments from either TWIST Bioscience (https://www.twistbioscience.com) or IDT (gBlocks, https://eu.idtdna.com/) resulted in the final experimental set of 17 sequences, with 4 from the expression bin of ~10 TPM, 6 from ~100 TPM and 7 from ~1000 TPM (Table S4).

### M5. Experimental strain construction

The *S. cerevisiae* strain S288C (ATCC no. 204508) was used as the base strain for all genetic engineering. Promoter (including 5’ UTR) and terminator (including 3’ UTR) DNA sequences were ordered as gene fragments from either TWIST Bioscience (https://www.twistbioscience.com) or IDT (https://eu.idtdna.com/). The exception were the RPL3 promoter and RPL3 terminator, for which fragments could not be synthesized due to sequence complexities, and were thus amplified from the genome with promoter_YOR063W_fwd, promoter_YOR063W_rev and terminator_YOR063W_fwd, terminator_YOR063W_rev primer pairs, respectively (Table S5). For the promoter-GFP-terminator constructs, the UBIMΔkGFP* version of the GFP gene from Houser et al. ^74^ was used (Table S6).

Integration of the promoter-GFP-terminator constructs into the genome at the XI-2 locus was done using the CRISPR/Cas9 plasmid (pCFB2312) and gRNA helper vectors (pCFB3044) from the EasyClone marker-free system ^75^. All transformation steps were performed according to the published manual, except that the repair fragment was provided as three fragments: the promoter with 90 bp overlap to the genome and 90 bp overlap to the GFP gene, the GFP gene, and the terminator with 90 bp overlap to the GFP gene and 90 bp overlap to the genome. The exception were the RPL3 promoter and terminator, which were amplified from the S288C genome with a shorter 40 bp overlap flanking the primers. For each fragment, the promoter, the GFP gene and the terminator were ligated together with a linearized pUC19 plasmid by Gibson assembly ^76^. The pUC19 vector was linearized by PCR with pUC19_fwd and pUC19_rev primer pair (Table S5), with 20 bp overlaps flanking the ends for the Gibson assembly. All plasmids were extracted using the Thermo Scientific GeneJET Plasmid Miniprep Kit and used as the templates for the promoter-GFP-terminator fragments with the L90 and R90 primer pair. To obtain strains with correctly integrated fragments at the XI-2 locus, colonies were verified with PCR using the 909 ^75^, GFP_rev and 910 ^75^, GFP_fwd primer pairs (Table S5) and the fragments were sequence-verified by Eurofins (https://www.eurofins.com/) after amplifying them with the L90, R90 primer pair.

For the mutagenesis experiment, designed promoter_RPL3 variants were ligated with GFP and the native terminator_RPL3 with the methods described above (Table S6). For the generative experiment, the different generated synthetic promoters and terminators corresponding to 17 whole constructs were ligated with GFP with the methods described above (Table S7).

### M6. RNA extraction and quantitative PCR

All yeast strains were cultured and monitored in a 48-well FlowerPlate (m2p-laboratories GmbH, Germany) at 30°C and 1200 rpm using a microbioreactor Biolector (m2p-laboratories GmbH, Germany). Cultures were started from a preculture grown overnight, at an OD600 of 0.03 in 1 mL minimal media with 2% glucose (Table S8). OD600 was monitored in real-time by the Biolector approximately every 20 min. After 15 h of cultivation, when the cells were in mid-exponential growth phase, the cells were collected and immediately used for RNA extraction with the QIAGEN RNeasy Mini Kit. For each batch of cultivation, the S288C wild type strain as well as the two integration strains with the POP6 and RPL3 regulatory regions were used as control groups. All cultivations were performed in biological triplicates.

cDNA was synthesized with QIAGEN QuantiTect Reverse Transcription Kit by adding 50 ng of total RNA to a final RT reaction volume of 20 μL. 1 μL of the cDNA was used as template with the Thermo Scientific DyNAmo Flash SYBR Green PCR Master Mix in a Mx3005P QPCR System (Agilent Technologies, USA). A 2 step qPCR protocol was used: 10 min initialization at 95^◦^C and 40 cycles of each: 30 s 95°C and 60 s 60°C. *S. cerevisiae* TAF10 ^77^ was selected as the reference gene and previously published primers were used, while primers for GFP were designed using IDT’s PrimerQuest tool (Table S5). Measurements were performed in separate batches due to the constraints of the measurement plate size to 96 wells. For each qPCR batch, samples from the S288C wild type strain as well as the two integration strains with the POP6 and RPL3 regulatory regions were included as the respective reference, low-expression and high-expression control groups. Each sample has technical duplicates. Cycle thresholds (Ct) of the reporter gene were normalized relative to the Ct value of TAF10 ^77^. The 2^−ΔΔCT^(avg. 2pddct) value was used as the indicator of the relative expression level of GFP for each construct ^78^, where the wild type strain was used as the reference (Table S2 and S4). The values were equalized across all qPCR batches based on the known TPM values of the native POP6 and RPL3 controls present in every batch, using a linear curve fit to infer the TPM values of each replicate of the generated constructs.

### M7. Data analysis and software

The performance of the generative model was monitored by measuring the sequence properties of the generated variants, including (i) sequence compositional validity, (ii) sequence similarity measures, (iii) predicted gene expression levels and (iv) known *cis*-regulatory grammar (Table S3), and by testing if they reflected the properties of natural sequences. DNA sequence homology was calculated with the *ratio* function in the python-Levenshtein package v0.12, equaling the Levenshtein (edit) distance divided by the length of the sequence. The Jaccard distance between two DNA sequences was defined as the intersection over union of sets of their unique k-mers of size 4. Nucleosome depletion was calculated using the R package nuCpos ^30,31^. Samples of 64 generated or natural test sequences were used per parameter except where stated otherwise. For statistical hypothesis testing, Scipy v1.1.0 was used with default settings. All tests were two-tailed except where stated otherwise. Python v3.6 (https://www.python.org) and R v3.6 (https://www.r-project.org) were used for computations.

## Author contributions

JZ and AZ conceptualized the project; JZ, ASM, NS, VV, MHC, DD and AZ designed the computational analysis; JZ, ASM and NS performed the computational analysis; JZ, XF, CS, VJ, CSB, VS, FD and AZ designed the experimental analysis; XF, CS and VJ performed the experimental analysis; JZ and AZ interpreted the results; JZ and AZ wrote the draft; All authors contributed to the final manuscript.

## Acknowledgements

We thank Sandra Viknander for technical discussions on deep neural networks. The study was supported by SciLifeLab funding and Swedish Research council (Vetenskapsrådet) starting grant 2019-05356. AZ was supported by Marius Jakulis Jason foundation.

