## Supplementary Information for "Supervised generative design of regulatory DNA for gene expression control"

#### Table of contents

Supplementary figures … p.2 - p.19

Supplementary tables … p.20 - p.38

Supplementary references … p.39 - p.40

### Supplementary figures

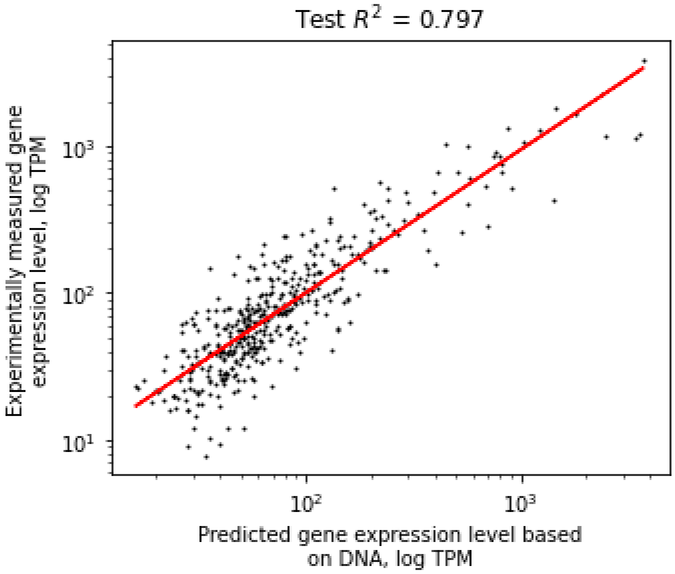


**Figure S1.** Performance of the predictive model of gene expression on the test dataset, trained on natural genomic sequences comprising whole gene regulatory structures of 1000 bps.

**A**

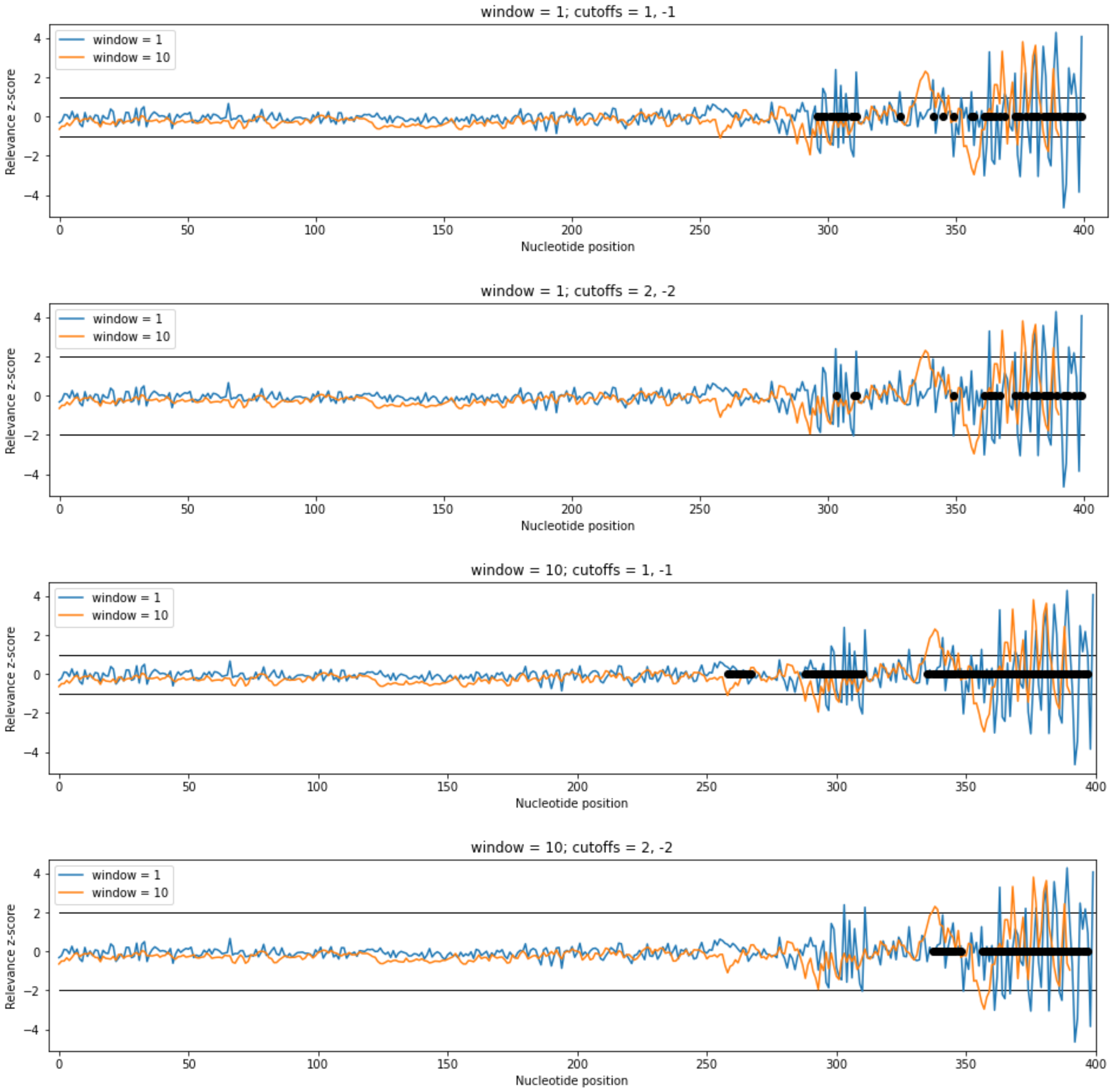
window = 1; cutoffs = 1, -1
window = 1
window = 10
Relevance z-score
Nucleotide position
window = 1; cutoffs = 2, -2
window = 1
window = 10
Relevance z-score
Nucleotide position
window = 10; cutoffs = 1, -1
window = 1
window = 10
Relevance z-score
Nucleotide position
window = 10; cutoffs = 2, -2
window = 1
window = 10
Relevance z-score
Nucleotide position

**B**

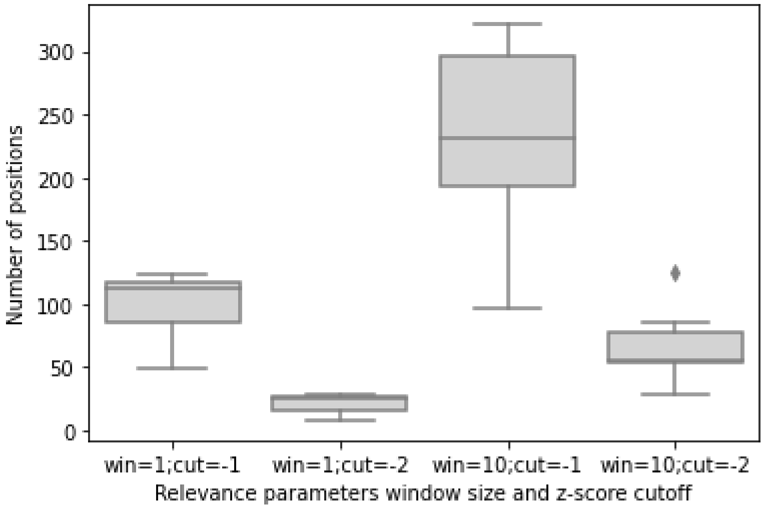


**Figure S2.** (A) Visualization of relevance profiles and relevant positions in the regulatory regions of the RPL3 gene and (B) total number of relevant positions, at different window size and z-score cutoff.

**A**

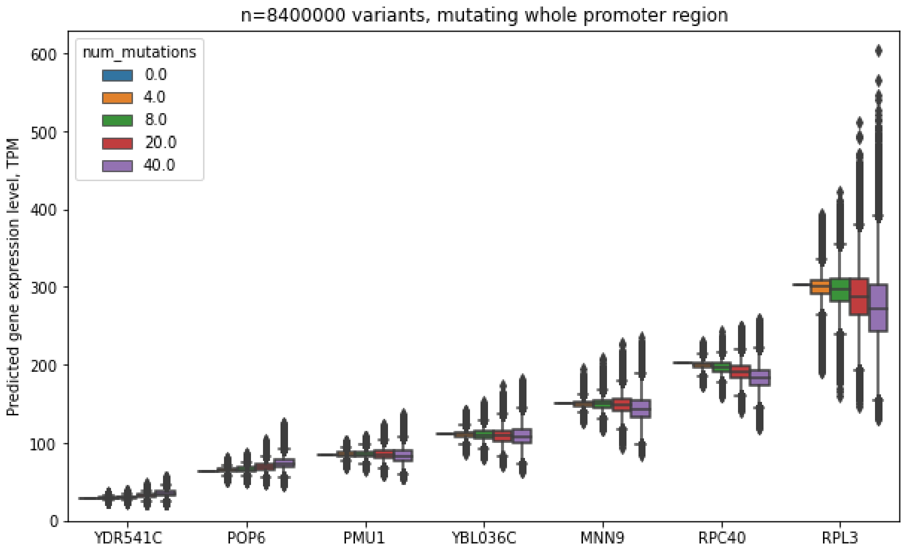


**B**

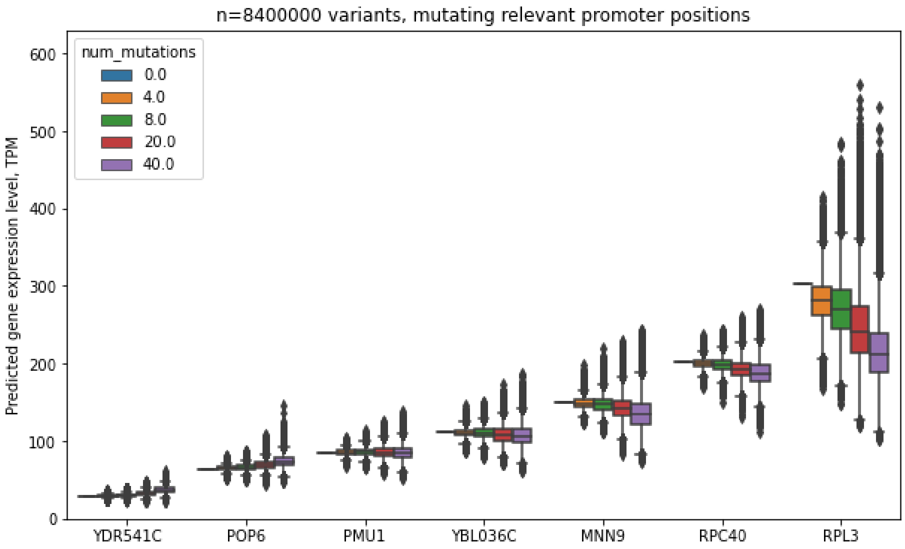


**Figure S3.** Predicted gene expression levels across sequence variants obtained by mutating (A) whole promoter regions and (B) only the most relevant positions as defined by querying the predictor's sensitivity.

**A**

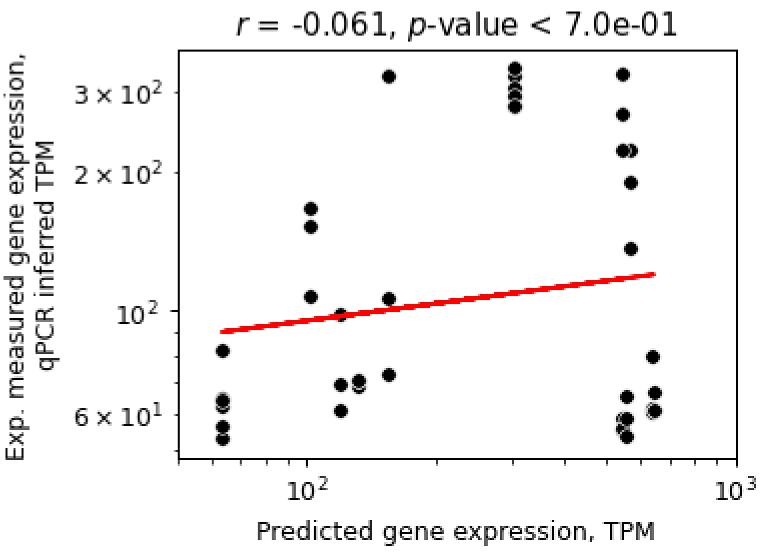


**B**

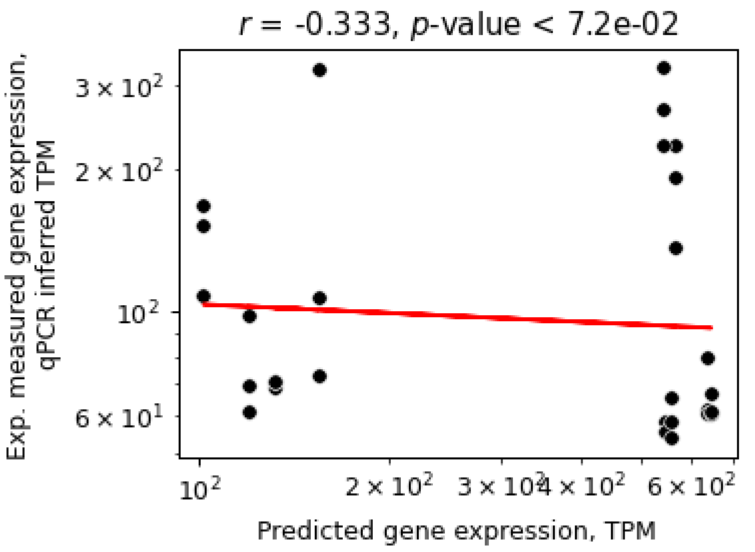


**Figure S4.** Correlation analysis of experimentally tested sequence variants from the mutational approach with (A) included controls and (B) only designed constructs.

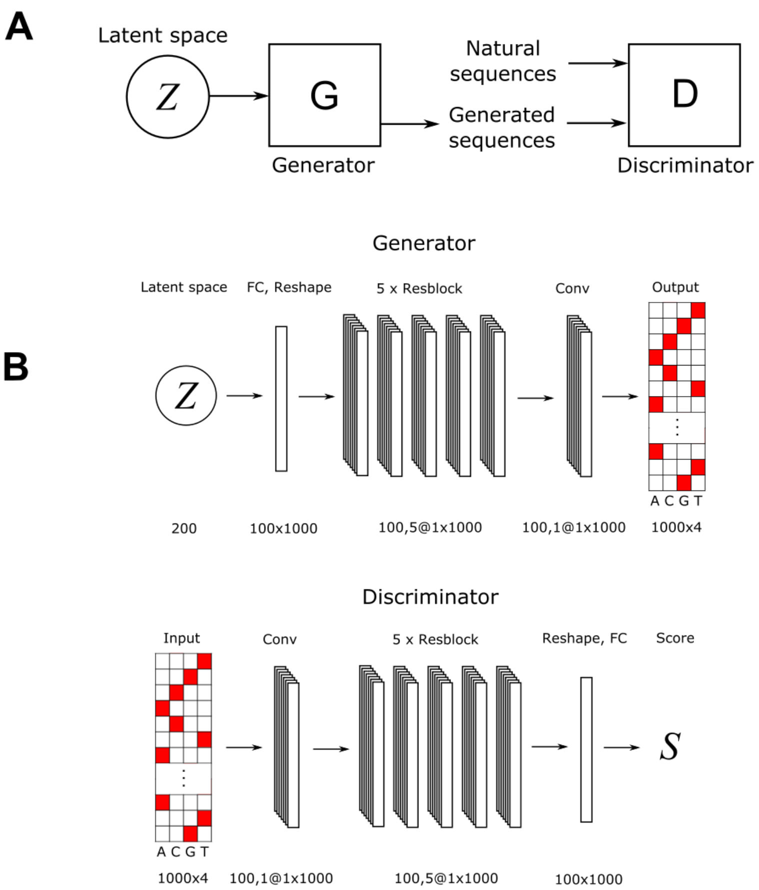


**Figure S5.** Overview of (A) generative adversarial network (GAN) approach and (B) its deep neural network architectures.

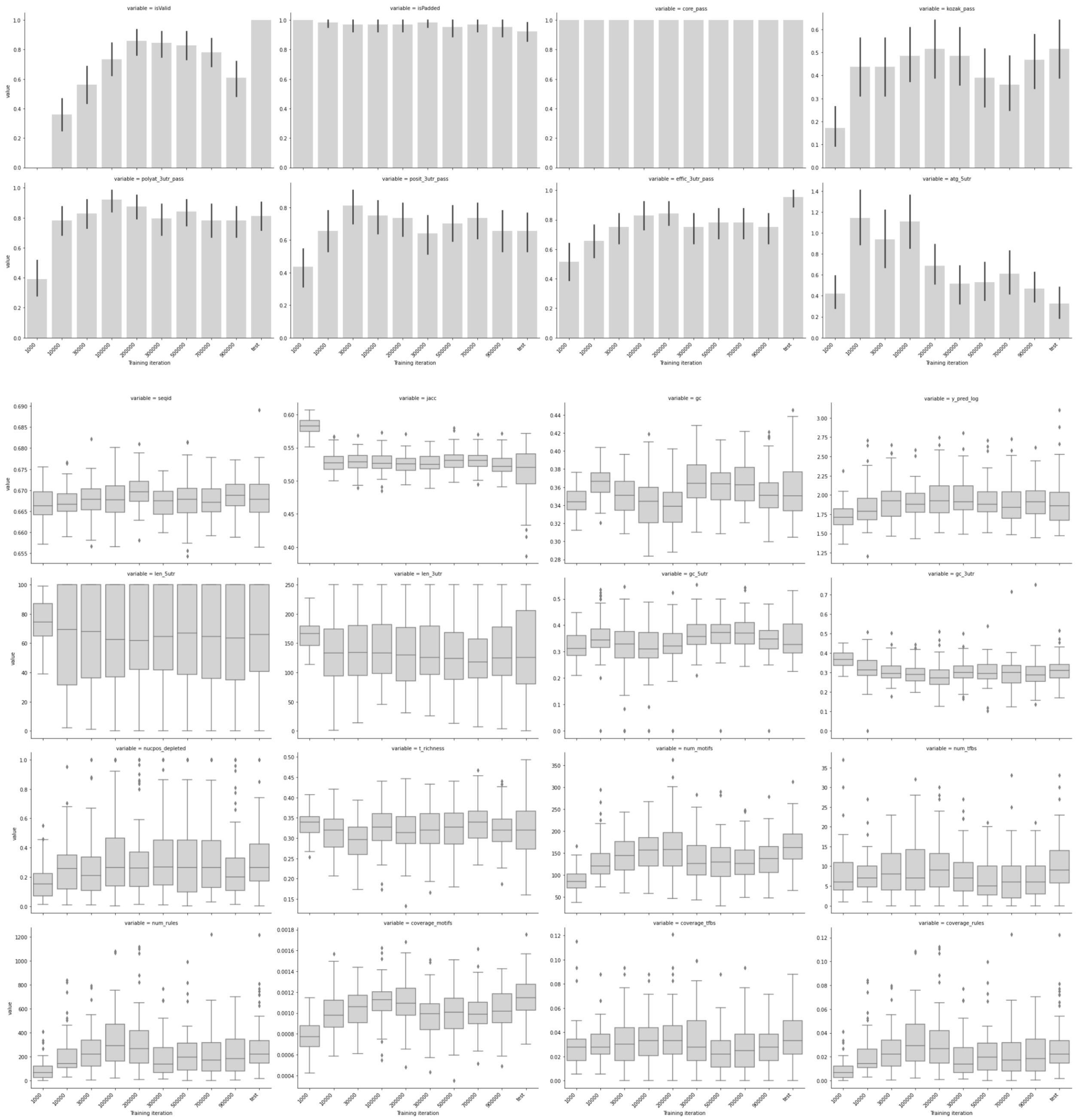


**Figure S6**. Computed sequence properties (see Table S3) of generated sequence variants sampled from the generative model at different numbers of training iterations.

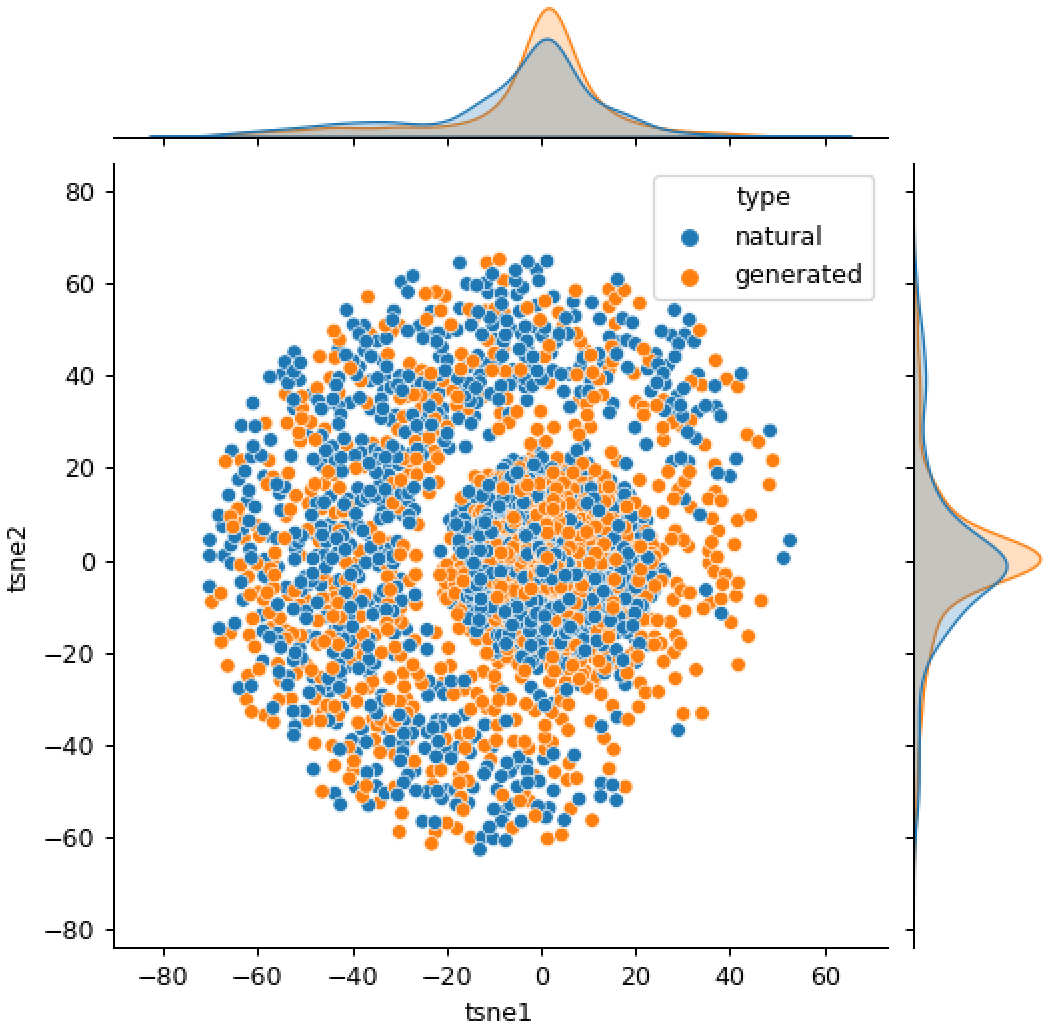


**Figure S7.** T-distributed stochastic neighbour embedding (t-SNE) dimensionality reduction (van der Maaten 2008) over the sequence identity distance matrix among equal amounts of combined natural and generated sequences.

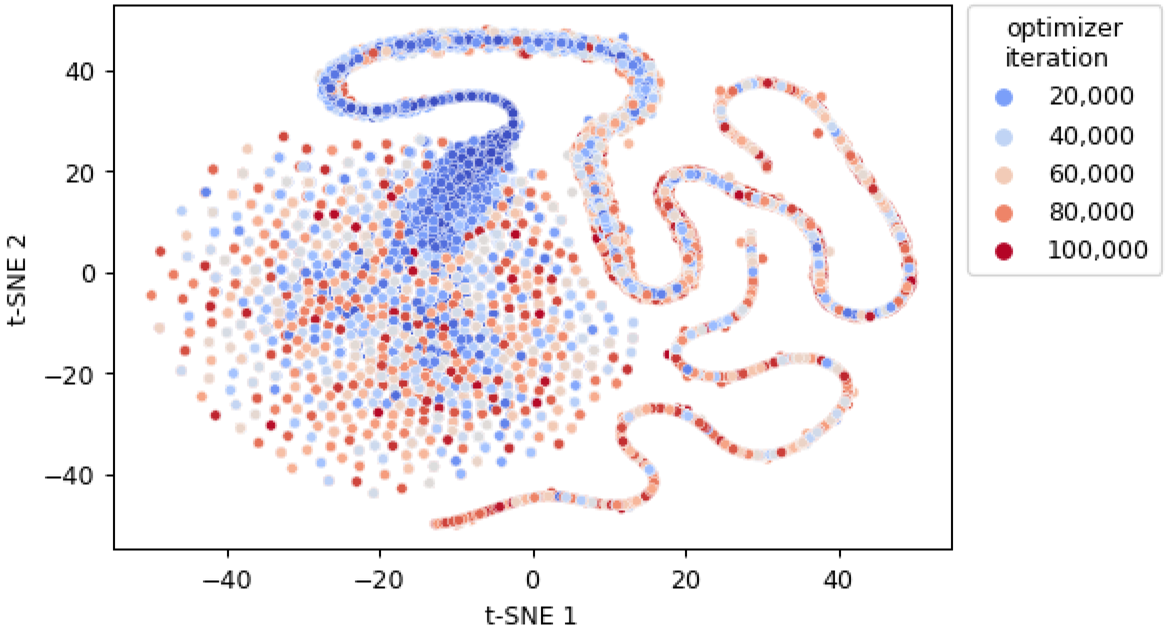


**Figure S8.** T-distributed stochastic neighbour embedding (t-SNE) dimensionality reduction (van der Maaten 2008) over the latent space vectors of generated sequence variants with the predictor-guided optimization approach, merging the results of both maximization and minimization of gene expression and marked by optimizer iterations.

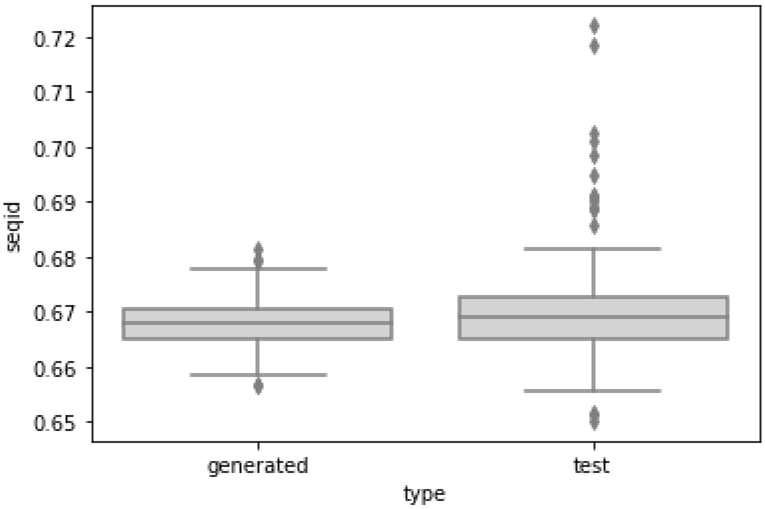


**Figure S9.** Sequence identity of generated and natural test set sequences versus their closest matching sequences in the training dataset.

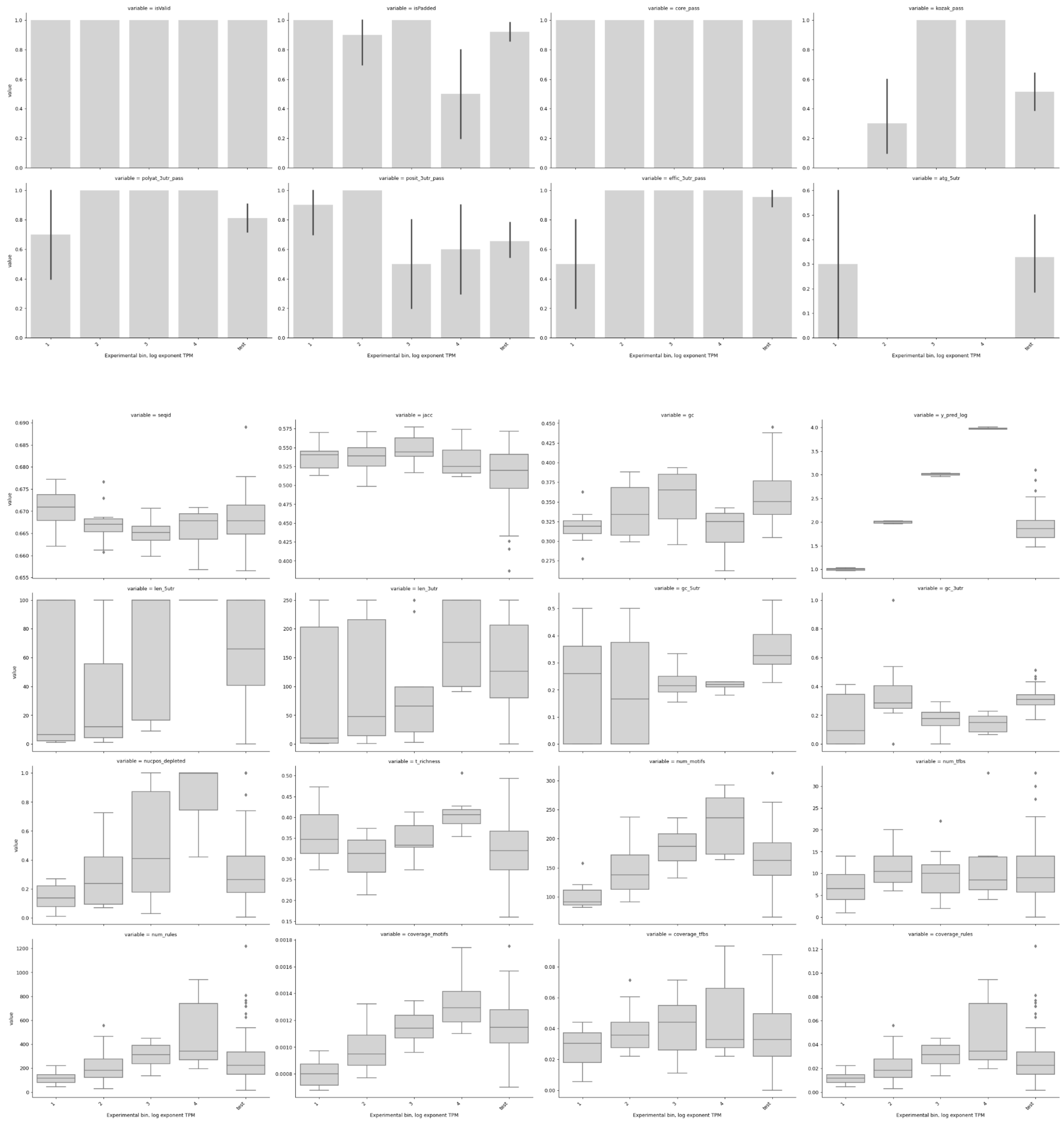


**Figure S10.** Computed sequence properties (see Table S3) of generated variants sampled across 4 orders of magnitude of expression levels (predicted TPM of 10, 100, 1000, 1000).

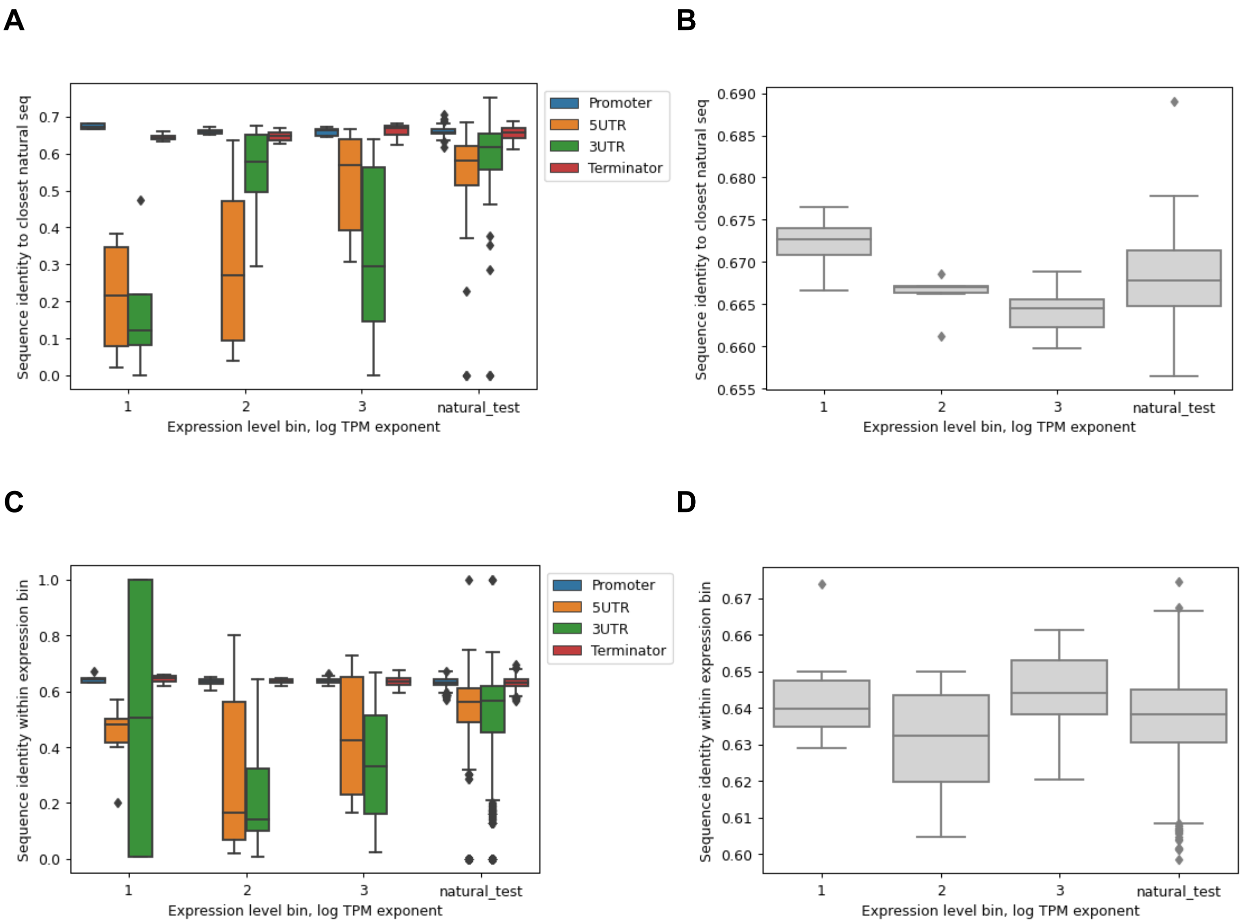


**Figure S11.** Sequence identity of generated and test sequences (A,B) to closest sequence in training dataset and (C,D) within experimental bins (defined by predicted expression levels of generated sequences), computed across (A,C) separate regions of the regulatory structures or (B,D) whole regulatory sequences.

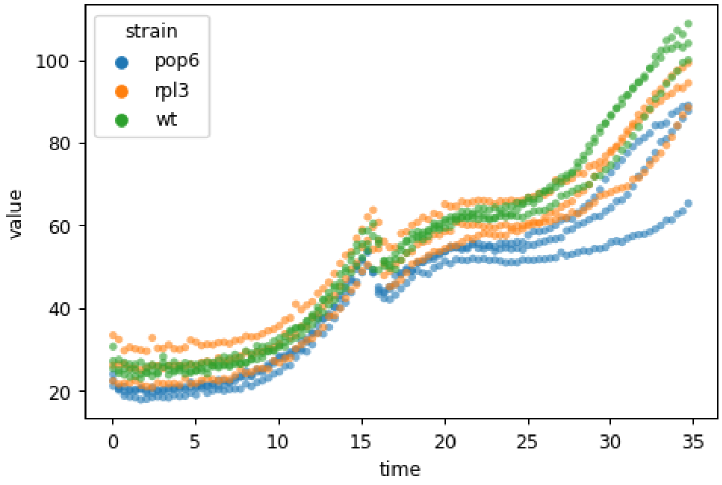


**Figure S12.** Biomass measurements using a bioreactor (Methods M5) showing that the GFP gene did not affect cell growth.

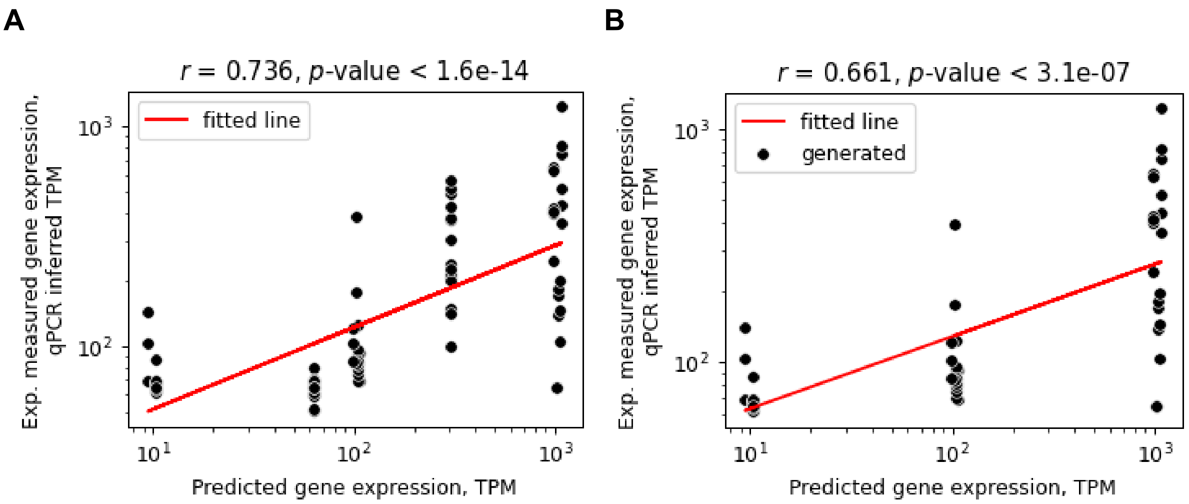


**Figure S13.** Correlation analysis across experimental bins of generated sequence variants (A) including all measured constructs (generated sequences and controls) or (B) including only generated sequences.

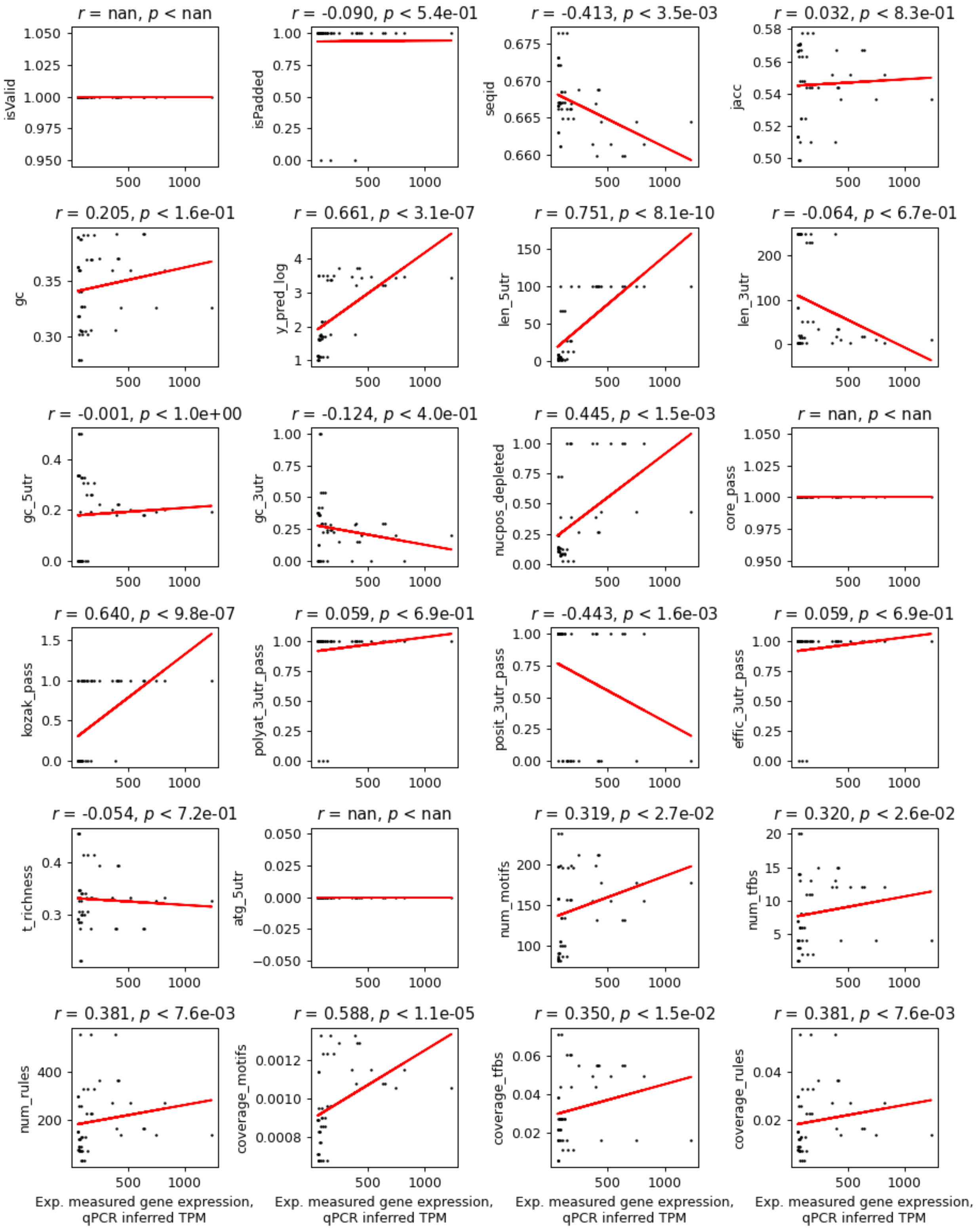


**Figure S14.** Correlation analysis between experimentally measured mRNA levels and predicted DNA sequence properties (see Table S3).

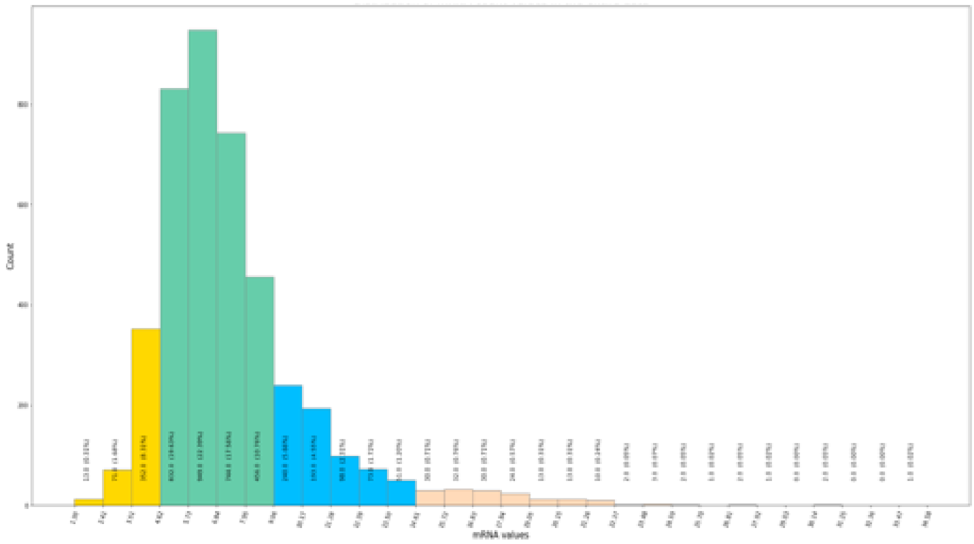


**Figure S15.** Binning the generative model training data across mRNA counts to perform data balancing.

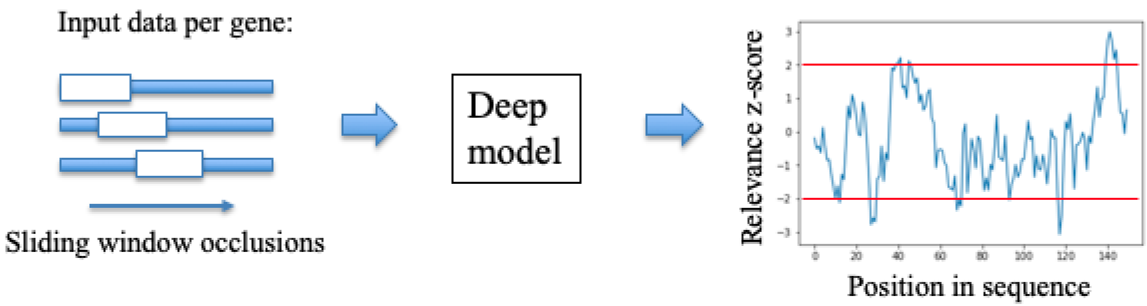


**Figure S16.** Schematic overview of the implemented DNA sequence occlusion-relevance approach (Zrimec et al. 2020; Ancona et al. 2017).

**A**

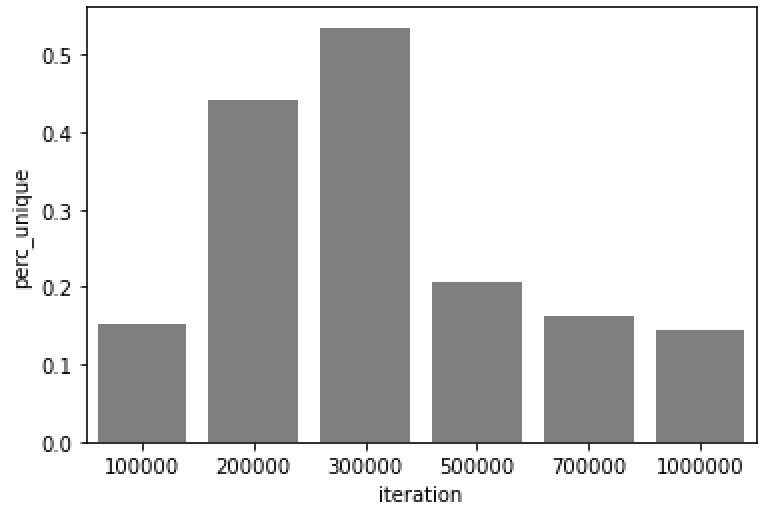


**B**

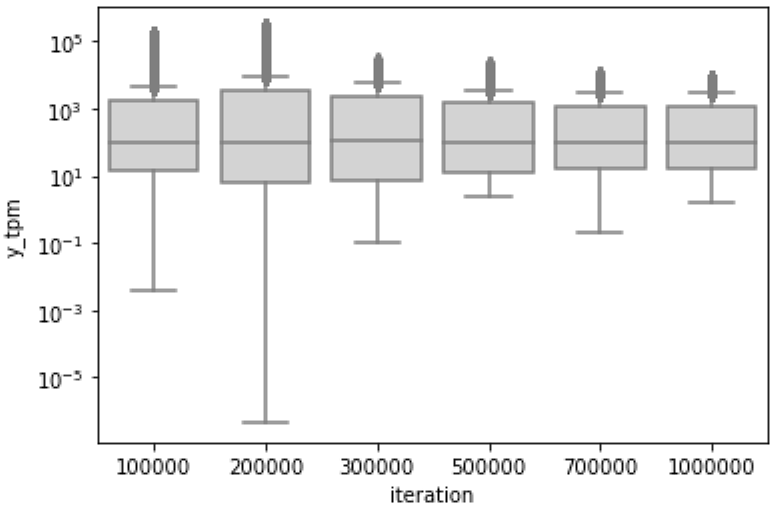


**Figure S17.** Generative model selection by comparing (A) the percentage of unique generated sequences and (B) the range of predicted gene expression levels, across generators obtained at different training iteration checkpoints and optimized for 100,000 iterations with the predictor-guided optimization procedure.

### Supplementary tables

**Table S1.** Number of sequence variants with >50% predicted change in gene expression levels obtained using the mutagenesis approach.

| region | window size | z-score cutoff | percent mutation | increase | decrease |
|---|---|---|---|---|---|
| relevant | 10 | 1 | 1 | 0 | 0 |
| | | | 2 | 0 | 0 |
| | | | 5 | 0.000013 | 0.00001 |
| | | | 10 | 0.000087 | 0.000667 |
| | 10 | 2 | 1 | 0 | 0 |
| | | | 2 | 0 | 0 |
| | | | 5 | 0.00001 | 0.000003 |
| | | | 10 | 0.000097 | 0.000107 |
| | 1 | 1 | 1 | 0 | 0 |
| | | | 2 | 0 | 0 |
| | | | 5 | 0.00004 | 0.000037 |
| | | | 10 | 0.00009 | 0.00158 |
| | 1 | 2 | 1 | 0 | 0 |
| | | | 2 | 0 | 0 |
| | | | 5 | 0 | 0.00002 |
| | | | 10 | 0 | 0.000027 |
| whole | / | / | 1 | 0 | 0 |
| | | | 2 | 0 | 0 |
| | | | 5 | 0.000007 | 0 |
| | | | 10 | 0.000077 | 0.000517 |

**Table S2.** Experimental validation (Methods M6) of designed sequence constructs with the mutational approach (Methods M3). Relevant positions were determined at window size 10 bp and z-score cutoff of 1.

| construct | bin | strategy | mutatated sequnce size, % | batch | predicted TPM | avg. 2pddct | qPCR inferred TPM |
|---|---|---|---|---|---|---|---|
| 3059399 | increase | relevant | 5 | 2 | 555.58 | 65.11 | 59.21 |
| 3080857 | increase | relevant | 5 | 2 | 645.09 | 86.43 | 62.81 |
| 3438152 | increase | relevant | 10 | 2 | 635.45 | 113.70 | 67.41 |
| 3480767 | increase | relevant | 10 | 2 | 544.09 | 48.97 | 56.48 |
| 8207535 | increase | whole | 10 | 1 | 541.38 | 1,689.55 | 273.57 |
| 8248101 | increase | whole | 10 | 1 | 564.84 | 975.18 | 183.82 |
| 3000527 | decrease | relevant | 5 | 2 | 119.88 | 165.75 | 76.20 |
| 3529331 | decrease | relevant | 10 | 1 | 101.67 | 644.39 | 142.25 |
| 7861215 | decrease | whole | 5 | 1 | 154.61 | 850.87 | 168.20 |
| 8172100 | decrease | whole | 10 | 1 | 131.41 | 71.11 | 70.22 |
| POP6 | ctrl low | / | / | 1 | 63.63 | 18.63 | 63.63 |
| POP6 | ctrl low | / | / | 2 | 63.63 | 91.29 | 63.63 |
| RPL3 | ctrl high | / | / | 1 | 303.20 | 1,925.34 | 303.20 |
| RPL3 | ctrl high | / | / | 2 | 303.20 | 1,509.69 | 303.20 |

**Table S3.** Verified DNA sequence properties. Properties with underlined variable names were used in the sequence selection procedure.

| Type | Variable name | Description |
|---|---|---|
| seq. similarity | seqid | Ratio of Levenshtein 'edit' distance to longer seq. length |
| seq. similarity | jacc | Jaccard distance |
| seq. composition | isPadded | Check if sequence contains Ns |
| seq. composition | isValid | Check if sequence contains Ns in correct positions (UTR borders only) |
| seq. composition | gc | GC-content |
| seq. composition | len_5utr | Length of 5' UTR region (Cheng et al. 2017; Neymotin, Ettorre, and Gresham 2016) |
| seq. composition | len_3utr | Length of 3' UTR region (Cheng et al. 2017; Neymotin, Ettorre, and Gresham 2016) |
| seq. composition | gc_5utr | GC-content of 5' UTR region (Cheng et al. 2017; Neymotin, Ettorre, and Gresham 2016) |
| seq. composition | gc_3utr | GC-content of 5' UTR region (Cheng et al. 2017; Neymotin, Ettorre, and Gresham 2016) |
| seq. composition | t_richness | T nucleotide richness in region from TSS to TATA box (region 0 to 150 bp from TSS analysed) (Lubliner, Keren, and Segal 2013) |
| seq. composition | homopolymer | Presence of homopolymers of length >9 bp, as they can limit synthesis |
| regulatory grammar | nucpos_depleted | Nucleosome depletion based on R package nuCpos (Xi et al. 2010; Kato, Shimizu, and Urano 2020) |
| regulatory grammar | core_pass | Presence of core promoter sequence 5'-TATAWAWR-3' (Basehoar, Zanton, and Pugh 2004; Lubliner, Keren, and Segal 2013) |
| regulatory grammar | kozak_pass | Presence of Kozak-like seq. in 5' UTR of 5-15 bp (Li et al. 2017; Nakagawa et al. 2008) |
| regulatory grammar | polyat_3utr_pass | Presence of poly-A/T seq. in 3' UTR or terminator (Curran et al. 2015; Zhao, Hyman, and Moore 1999; van Helden, del Olmo, and Pérez-Ortín 2000) |
| regulatory grammar | posit_3utr_pass | Presence of positioning element 5'-AAWAAA-3 in 3' UTR or terminator (Guo and Sherman 1996; Curran et al. 2015) |
| regulatory grammar | effic_3utr_pass | Presence of efficiency element 5'-TATDTA-3 in 3 'UTR or terminator (Guo and Sherman 1996; Shalem et al. 2015) |
| regulatory grammar | atg_5utr | Presence of upstream ATGs in 5' UTR (Dvir et al. 2013; Cheng et al. 2017) |
| regulatory grammar | num_motifs | Overall num. relevant sequence motifs identified from reference study (Zrimec et al. 2020) |
| regulatory grammar | num_tfbs | Overall num. TFBS from the Jaspar and Yeastract databases (Khan et al. 2018; Teixeira et al. 2018) |
| regulatory grammar | num_rules | Overall num. co-occurring motifs identified from reference study (Zrimec et al. 2020) |
| regulatory grammar | coverage_motifs | Coverage of relevant sequence motifs identified from reference study (Zrimec et al. 2020) |
| regulatory grammar | coverage_tfbs | Coverage of TFBS from the Jaspar and Yeastract databases (Khan et al. 2018; Teixeira et al. 2018) |
| regulatory grammar | coverage_rules | Coverage of co-occurring motifs identified from reference study (Zrimec et al. 2020) |
| model predictions | y_pred | Predicted gene expression level (Zrimec et al. 2020) |

**Table S4.** Experimental validation (Methods M6) of designed sequence constructs with the generative approach (Methods M4).

| construct | bin | batch | predicted TPM | avg. 2pddct | qPCR inferred TPM |
|---|---|---|---|---|---|
| 5 | gen ~10 | 5 | 10.33 | 12.65 | 63.12 |
| 6 | gen ~10 | 5 | 10.41 | 28.35 | 66.47 |
| 7 | gen ~10 | 5 | 10.35 | 64.67 | 74.22 |
| 10 | gen ~10 | 4 | 9.45 | 13.95 | 104.65 |
| 15 | gen ~100 | 2 | 104.04 | 51.69 | 83.33 |
| 17 | gen ~100 | 4 | 97.94 | 13.64 | 102.90 |
| 18 | gen ~100 | 2 | 103.65 | 36.74 | 74.23 |
| 19 | gen ~100 | 2 | 103.86 | 85.74 | 104.07 |
| 20 | gen ~100 | 2 | 106.48 | 47.35 | 80.69 |
| 23 | gen ~100 | 4 | 103.33 | 33.72 | 216.87 |
| 26 | gen ~1000 | 2 | 1,027.04 | 21.78 | 65.12 |
| 27 | gen ~1000 | 3 | 1,055.05 | 9.38 | 149.07 |
| 28 | gen ~1000 | 1 | 1,046.14 | 9.70 | 163.82 |
| 29 | gen ~1000 | 1 | 989.01 | 39.54 | 558.62 |
| 31 | gen ~1000 | 1 | 1,076.07 | 58.27 | 806.45 |
| 33 | gen ~1000 | 4 | 1,077.47 | 95.58 | 567.97 |
| 34 | gen ~1000 | 3 | 985.57 | 27.30 | 358.57 |
| POP6 | ctrl low | 1 | 63.63 | 2.13 | 63.63 |
| POP6 | ctrl low | 2 | 63.63 | 19.33 | 63.63 |
| POP6 | ctrl low | 3 | 63.63 | 2.08 | 63.63 |
| POP6 | ctrl low | 4 | 63.63 | 6.72 | 63.63 |
| POP6 | ctrl low | 5 | 63.63 | 15.05 | 63.63 |
| RPL3 | ctrl high | 1 | 303.20 | 20.24 | 303.20 |
| RPL3 | ctrl high | 2 | 303.20 | 412.72 | 303.20 |
| RPL3 | ctrl high | 3 | 303.20 | 22.56 | 303.20 |
| RPL3 | ctrl high | 4 | 303.20 | 48.93 | 303.20 |
| RPL3 | ctrl high | 5 | 303.20 | 1,137.72 | 303.20 |

**Table S5.** List of PCR primers.

| primer name | sequence |
|---|---|
| L90 | TTTAGCACGCGGGGTGTAA |
| R90 | TTTCATCAAGAGAAGAAACACG |
| promoter_YOR063W_fwd | TAGGCAAAAGCCAAGGAGCGTTTGCCATGAACTTCCACAATTATTAAA TTCAGTGGTAATGCAA |
| promoter_YOR063W_rev | TTATGGTTTTACCGGTCAAAGTCTTGACGAAAATCTGCATTGATTGATT GTTGTAGTAACTGTG |
| terminator_YOR063W_fwd | TGCTGGGATTACACATGGCATGGATGAACTATACAAATAGAGAAGTTT TGTTAGAAAATAAATCATTTTT |
| terminator_YOR063W_rev | ACATCTAAACTTTTTAATATCTGAAAGCGCTAGTCGTGTGGGCTTGTCC CTTCGAGTG |
| pUC19_fwd | TGTTTCTTCTCTTGATGAAAGGGTACCGAGCTCGAATTC |
| pUC19_rev | GTTACACCCCGCGTGCTAAAGGGGATCCTCTAGAGTCG |
| 909 | GTTTGTAGTTGGCGGTGGAG |
| 910 | GAGACAAGATGGGGCAAGAC |
| GFP_rev | AATTTTCGACTTAACGTTGTCG |
| GFP_fwd | GATCCCAACGAAAAGAGAG |
| GFP_qPCR_fwd | GTCACTACTCTCACTTATGGTGTTC |
| GFP_qPCR_rev | GTGTCTTGTAGTTCCCGTCATC |
| TAF10_qPCR _fwd | ATATTCCAGGATCAGGTCTTCCGTAGC |
| TAF10_qPCR _rev | GTAGTCTTCTCATTCTGTTGATGTTGTTGTTG |

**Table S6.** List of DNA fragments used in constructs for the mutagenesis experiment.

| construct | sequence |
|---|---|
| *UBIMΔkGFP** | ATGCAGATTTTCGTCAAGACTTTGACCGGTAAAACCATAACATTGGAAGTTG<br>AATCTTCCGATACCATCGACAACGTTAAGTCGAAAATTCAAGACAAGGAAGG<br>TATCCCTCCAGATCAACAAAGATTGATCTTTGCCGGTAAGCAGCTAGAAGAC<br>GGTAGAACGCTGTCTGATTACAACATTCAGAAGGAGTCCACCTTACATCTTG<br>TGCTAAGGCTAAGAGGTGGTATGCACGGATCCGGAGCTTGGCTGTTGCCCG<br>TCTCACTGGTGAAAAGAAAAACCACCCTGGCGCCCAATACGAGTAAAGGAG<br>AAGAACTTTTCACTGGAGTTGTCCCAATTCTTGTTGAATTAGATGGTGATGTT<br>AATGGGCACAAATTTTCTGTCAGTGGAGAGGGTGAAGGTGATGCAACATAC<br>GGAAAACTTACCCTTAAATTTATTTGCACTACTGGAAAACTACCTGTTCCATG<br>GCCAACACTTGTCACTACTCTCACTTATGGTGTTCAATGCTTTTCAAGATACC<br>CAGATCACATGAAACAGCATGACTTTTTCAAGAGTGCCATGCCCGAAGGTTA<br>TGTACAGGAAAGAACTATATTTTTCAAAGATGACGGGAACTACAAGACACGT<br>GCTGAAGTCAAGTTTGAAGGTGATACCCTTGTTAATAGAATCGAGTTAAAAG<br>GTATTGATTTTAAAGAAGATGGAAACATTCTTGGACACAAATTGGAATACAAC<br>TATAACTCACACAATGTATACATCATGGCAGACAAACAAAAGAATGGAATCAA<br>AGCTAACTTCAAAATTAGACACAACATTGAAGATGGAAGCGTTCAACTAGCA<br>GACCATTATCAACAAAATACTCCAATTGGCGATGGCCCTGTCCTTTTACCAGA<br>CAACCATTACCTGTCCACACAATCTGCCCTTTCGAAAGATCCCAACGAAAAG<br>AGAGACCACATGGTCCTTCTTGAGTTTGTAACAGCTGCTGGGATTACACATG<br>GCATGGATGAACTATACAAATAG |
| promoter_POP6 | TTTAGCACGCGGGGTGTAACTCAACAGAAAAATGTGCCATAGAACAAGACTA<br>GGCAAAAGCCAAGGAGCGTTTGCCATGAACTTCCACAATCTCTTGATTATGT<br>CATATGAAAGGTCCAGTGGGACTTGCTTTTGTTGCAGCACCTTTGCTAATGA<br>ATGAAAGGCACATAGTGACTGCTTAAAAATGCAGGAACTTAAATTATTCCGAA<br>TGGTATTTTGTCTCACATATATTGTCCCATACTGTGCCAGGATCCCGGCTTTA<br>CCCAGTATCATCATTGTACCGTTACCAATTCTCCTCGTATATCACGGTTAGTTT<br>TTAAACCTCGGGGTGACGTTTACTATTGGCGTACTAATATATTCTTATTTTCTT<br>TTCTTTTTTGTTGGCAGTTTCAAGCAACACATGTACTGGATAACCAACCCCC<br>GCACGCTCTTGGAAAAAATTGAGAAGGCATCGGACACTTGCTGATGAGTATT<br>TCGAAAAATTCCATGAAAGATGAGGCCAAGATTGTTTGGAAGAGATTGAAAA<br>GAAGAAGAAGAAAAAAAGATAAAAGCAAATCAAAAATGCAGATTTTCGTCAA<br>GACTTTGACCGGTAAAACCATAACATTGGAAGTTGAATCTTCCGATACCATCG<br>ACAACGTTAAGTCGAAAATT |
| terminator_POP6 | GATCCCAACGAAAAGAGAGACCACATGGTCCTTCTTGAGTTTGTAACAGCT<br>GCTGGGATTACACATGGCATGGATGAACTATACAAATAGAATCGACCAGCTC<br>TTTTAGCATCCATAGCTACTTCTTGCATTTGTACTTTATAATATAAAGCATTTTT<br>AGAAGGCTTAATCGATATCAGAACTACCAATTGTTCCTACTACAAGAAGTATG<br>TGTCATTGAATGAAAGAAAAAAGGATGCATGGAAATAGCACAACTTTATTTAT<br>TTTCTTCCCTTTGGAAATCGGAAATTCAATGATATGCCCTCTGATATAATAGCA<br>AAGACATGATCGTTAATTTAGGCCTCGCTTTGAATCCCACACGACTAGCGCT<br>TTCAGATATTAAAAAGTTTAGATGTAGGTTTTAGCGGTAACAGTTATATAAATC<br>GTGTTTCTTCTCTTGATGAAA |
| promoter_RPL3 | TAGGCAAAAGCCAAGGAGCGTTTGCCATGAACTTCCACAATTATTAAATTCA<br>GTGGTAATGCAACAGCAAGAGGAAAGGTGGAGGGATTAACGCATTTCAGAC<br>AGCTTTATAGGGGGAAAGAAAGCACTCGCAAACTTGCTGCCTGTTCGCAGT<br>CATTGGTTGCAAAAACTAAACTCTACTCACGCACACTGGAATGAATGGCAAT<br>ATTCTTTTTTAGGTTAACCGGCCGGACAGTAATATAGTAATCGTTTTGTACGTT<br>TTTCAAGAAGCGACGCACAACTGTTTTCCATTTTTTTTTTTTTTTTTTCAGTGA<br>TCATCGTCCATGAAAAAAATTTTTCATTTGTCTCTTTCGTGCTTCCTGGATATA<br>TAAAATACGATTTATTTAGTTGTCTTTGTCAATCCTCATCTTTCTTTACTCATTA<br>TTTCATTTCGGTTTTGTTCATCTCTAGAACAACACAGTTACTACAACAATCAAT<br>CAATGCAGATTTTCGTCAAGACTTTGACCGGTAAAACCATAA |

| | |
|---|---|
| terminator_RPL3 | TGCTGGGATTACACATGGCATGGATGAACTATACAAATAGAGAAGTTTTGTTAGAAAATAAATCATTTTTTAATTGAGCATTCTTATTCCTATTTTATTTAAATAGTTTTATGTATTGTTAGCTACATACAACAGTTTAAATCAAATTTTCTTTTTCCCAAAGTCCAAAATGGAGGTTTATTTTGATGACCCGCATGCGATTATGTTTTGAAAGTATAAGACTACATACATGTACATATATTTAAACATGTAAACCCGTCCATTATATTGCTTACTTTCTTCTTTTTTGCCGTTTTGACTTGGACCTCTGGTTTGCTATTTCCTTACAATCTTTGCTACAATACCATTTGCCCTTGGGAGCTTGTTTCAGGCCTACGCAACCATAATGGAACCACTCGAAGGGACAAGCCCACACGACTAGCGCTTTCAGATATTAAAAAGTTTAGATGT |
| promRPL3_variant3000527 | TTTAGCACGCGGGGTGTAACTCAACAGAAAAATGTGCCATAGAACAAGACTAGGCAAAAGCCAAGGAGCGTTTGCCATGAACTTCCACAATTATTAAATTCAGTGGTAATGCAACAGCAAGAGGAAAGGTGGAGGGATTAACGCATTTCAGACAGCTTTATAGGGGGAAAGAAAGCACTCGCAAACTTGCTGCCTGTTCGCAGTCATTGGTTGCAAAAACTAAACTCTACTCACGCACACTGGAATGAATGGCAATATTCTTTTTTAGGTTAACCGGCCGGACAGTAATATAGTAATCGTTTTGTACGTTTTTCAAGAAGCGACGCACAACTGTTTTCCATTTTTTTTTTTTTAATTTCAGTGATCATCGTCCATGAATAAAATTTTTGATTTGTCTTATTCGTGCTTCCTGGATATATAAAATACGATTACTTTAGTTATCTTCGTTAATCCCCATCTATCTATACTCATCATCACACTTCGGATGTGTTCATCTCTAGAACAACACAGTTACTACAACAATCAATCAATGCAGATTTTCGTCAAGACTTTGACCGGTAAAACCATAACATTGGAAGTTGAATCTTCCGATACCATCGACAACGTTAAGTCGAAAATT |
| promRPL3_variant3059399 | TTTAGCACGCGGGGTGTAACTCAACAGAAAAATGTGCCATAGAACAAGACTAGGCAAAAGCCAAGGAGCGTTTGCCATGAACTTCCACAATTATTAAATTCAGTGGTAATGCAACAGCAAGAGGAAAGGTGGAGGGATTAACGCATTTCAGACAGCTTTATAGGGGGAAAGAAAGCACTCGCAAACTTGCTGCCTGTTCGCAGTCATTGGTTGCAAAAACTAAACTCTACTCACGCACACTGGAATGAATGGCAATATTCTTTTTTAGGTTAACCGGCCGGACAGTAATATAGTAATCGTTTTGTACGTTTTTCAAGAAGCGACGCACAACTGTTTTCCATTTTTTTTTTCGTTTTTTCAGTGATCATCGTCCATGAAAAAAATCTGTCATTGCACTCTTTCGTGCTTCCTGGATATATAAAATACGAGTTATTTAGTTGTATTTGTCAAGTTTCTTCCTTACTTTCTTATTATTTTATTTCCGTTTTGTTCATCTCTAGAACAACACAGTTACTACAACAATCAATCAATGCAGATTTTCGTCAAGACTTTGACCGGTAAAACCATAACATTGGAAGTTGAATCTTCCGATACCATCGACAACGTTAAGTCGAAAATT |
| promRPL3_variant3080857 | TTTAGCACGCGGGGTGTAACTCAACAGAAAAATGTGCCATAGAACAAGACTAGGCAAAAGCCAAGGAGCGTTTGCCATGAACTTCCACAATTATTAAATTCAGTGGTAATGCAACAGCAAGAGGAAAGGTGGAGGGATTAACGCATTTCAGACAGCTTTATAGGGGGAAAGAAAGCACTCGCAAACTTGCTGCCTGTTCGCAGTCATTGGTTGCAAAAACTAAACTCTACTCACGCACACTGGAATGAATGGCAATATTCTTTTTTAGGTTAACCGGCCGGACAGTAATATAGTAATCGTTTTGTACGTTTTTCAAGAAGCGACGCACAACTGTTTTCCATTTTTTTTTTTTTGTTGGCAGTGATCATCGTCCATGAAAAAAGTTTTTCGTAAGTATCTTGCGTGCTTCCTGGATATATAAAATACGTTTTATTTAGGTGGCTATGTCAATCCTCTTCCTTCTTTCCTCGTTCTTTCTTTTCCGTTTTGTTCATCTCTAGAACAACACAGTTACTACAACAATCAATCAATGCAGATTTTCGTCAAGACTTTGACCGGTAAAACCATAACATTGGAAGTTGAATCTTCCGATACCATCGACAACGTTAAGTCGAAAATT |

| | |
|---|---|
| promRPL3_variant3438152 | TTTAGCACGCGGGGTGTAACTCAACAGAAAAATGTGCCATAGAACAAGACTA<br>GGCAAAAGCCAAGGAGCGTTTGCCATGAACTTCCACAATTATTAAATTCAGT<br>GGTAATGCAACAGCAAGAGGAAAGGTGGAGGGATTAACGCATTTCAGACAG<br>CTTTATAGGGGGAAAGAAAGCACTCGCAAACTTGCTGCCTGTTCGCAGTCA<br>TTGGTTGCAAAAACTAAACTCTACTCACGCACACTGGAATGAATGGCAATATT<br>CTTTTTTAGGTTAACCGGCCGGACAGTAATATAGTAATCGTTTTGTACGTTTTT<br>CAAGAAGCGACGCACAACTGTTTTCCATTTTTTTTTGCTTCTTCACAGTGATC<br>ATCGTCCATGAATAACCCTGAGCATGCGTCGTGTTCGTGCTTCCTGGATATAT<br>AAAATACTAATAGGTGAGTTTTATATGACAAGCTTAATTGTTCTTTTCTCTTTGT<br>TTCATTGATTTATTGTTCATCTCTAGAACAACACAGTTACTACAACAATCAATC<br>AATGCAGATTTTCGTCAAGACTTTGACCGGTAAAACCATAACATTGGAAGTT<br>GAATCTTCCGATACCATCGACAACGTTAAGTCGAAAATT |
| promRPL3_variant3480767 | TTTAGCACGCGGGGTGTAACTCAACAGAAAAATGTGCCATAGAACAAGACTA<br>GGCAAAAGCCAAGGAGCGTTTGCCATGAACTTCCACAATTATTAAATTCAGT<br>GGTAATGCAACAGCAAGAGGAAAGGTGGAGGGATTAACGCATTTCAGACAG<br>CTTTATAGGGGGAAAGAAAGCACTCGCAAACTTGCTGCCTGTTCGCAGTCA<br>TTGGTTGCAAAAACTAAACTCTACTCACGCACACTGGAATGAATGGCAATATT<br>CTTTTTTAGGTTAACCGGCCGGACAGTAATATAGTAATCGTTTTGTACGTTTTT<br>CAAGAAGCGACGCACAACTGTTTTCCATTTTTTTTTTTCAATAGTCAGTGATC<br>ATCGTCCATGAAGGACATCAGCCAGGTGGCTCTGTCGTGCTTCCTGGATATA<br>TAAAATATCTTTTCTTGAGTTGAATTTGTACATTGTTTTCGATCGTTACTTATTC<br>CTTCCGTTCTTATTTGTTCATCTCTAGAACAACACAGTTACTACAACAATCAAT<br>CAATGCAGATTTTCGTCAAGACTTTGACCGGTAAAACCATAACATTGGAAGTT<br>GAATCTTCCGATACCATCGACAACGTTAAGTCGAAAATT |
| promRPL3_variant3529331 | TTTAGCACGCGGGGTGTAACTCAACAGAAAAATGTGCCATAGAACAAGACTA<br>GGCAAAAGCCAAGGAGCGTTTGCCATGAACTTCCACAATTATTAAATTCAGT<br>GGTAATGCAACAGCAAGAGGAAAGGTGGAGGGATTAACGCATTTCAGACAG<br>CTTTATAGGGGGAAAGAAAGCACTCGCAAACTTGCTGCCTGTTCGCAGTCA<br>TTGGTTGCAAAAACTAAACTCTACTCACGCACACTGGAATGAATGGCAATATT<br>CTTTTTTAGGTTAACCGGCCGGACAGTAATATAGTAATCGTTTTGTACGTTTTT<br>CAAGAAGCGACGCACAACTGTTTTCCATTTTTTTTTGCTTTTTCTCAGTGATC<br>ATCGTCCATGAAACTATTTTTTTTATTGATTTTTTCGTGCTTCCTGGATATATAA<br>AATAGGCTCAATTTAGTTTATTTGGACAATCAGCATCGTAATCAGCTCCTATAA<br>AAATAGCGGAATTGTTCATCTCTAGAACAACACAGTTACTACAACAATCAATC<br>AATGCAGATTTTCGTCAAGACTTTGACCGGTAAAACCATAACATTGGAAGTT<br>GAATCTTCCGATACCATCGACAACGTTAAGTCGAAAATT |
| promRPL3_variant7861215 | TTTAGCACGCGGGGTGTAACTCAACAGAAAAATGTGCCATAGAACAAGACTA<br>GGCAAAAGCCAAGGAGCGTTTGCCATGAACTTCCACAATTATTAAATTCAGT<br>GGTAATGCAACAGCAAGAGGAAAGATGGGGCGATTAACGCATTTCAGACAG<br>CTTTATAGGGGGAAAGAAAGCACTCGCAAACTTGCTGCCTGTTCACAGTCAT<br>TGGTTGCAAAAACTAAACTCTACTCTCGCTCACTGGAATGAATGGCAATATTC<br>TTTTTTAGGTTAACCGGCAGGATAGTAATATAGTAATCGTTTTGTACGTTTTTC<br>AAGAAGCGACGCACAACTGTTTTCCATTATTTTTTTGTTTTTTTCAGTGATCAT<br>CGTCCATTAAAAAAATTTTTCATTTGTCTCTTTAGTGCTTCCTGGATATATAAAA<br>TACGATTGATTTAATTGTCTTTGACAATCCTCATCTTCCTTTAATCATCGATTCA<br>TTTCGGTTTTGTTCATCTCTAGAACAACACAGTTACTACAACAATCAATCAAT<br>GCAGATTTTCGTCAAGACTTTGACCGGTAAAACCATAACATTGGAAGTTGAA<br>TCTTCCGATACCATCGACAACGTTAAGTCGAAAATT |

| | |
|---|---|
| promRPL3_variant8172100 | TTTAGCACGCGGGGTGTAACTCAACAGAAAAATGTGCCATAGAACAAGACTA<br>GGCAAAAGCCAAGGAGCGTTTGCCATGAACTTCCACAATGATTAAATTCAGG<br>GGTAATGCAGCAGCAAGAGGAAAGGTGGGGCGATTAACCCATTTCAGACAG<br>CCTTATAGGGGGAAGGAAAGCACTCGCAATCTTGCTGCCTTTTCGCTGTAAT<br>TGGTTGCAAAAACTAGACTCTACTCACGCACACTGGAATGAATGTGAATATT<br>CTTTTTTAGGTCAACAGGCCGGACAGTAAAATAGTAATTGTTTTGGACGTTTT<br>TCAAGAAGCGACGCACAACTGTTTTCCATTTTTTTCTTTTTTTTTAAAGTGAT<br>CATACTCCATGAAAAAAATGTTTAATTTGTCTCTTTCGTGCTACCTGGATATAT<br>AAAATACGATCACTTTAGTTTTCTTTGCCCATCCTCATCTATCACTTCTCATTAT<br>ATCATTTCGGTATTGTTCATCTCTAGAACAACACAGTTACTACAACAATCAATC<br>AATGCAGATTTTCGTCAAGACTTTGACCGGTAAAACCATAACATTGGAAGTT<br>GAATCTTCCGATACCATCGACAACGTTAAGTCGAAAATT |
| promRPL3_variant8207535 | TTTAGCACGCGGGGTGTAACTCAACAGAAAAATGTGCCATAGAACAAGACTA<br>GGCAAAAGCCAAGGAGCGTTTGCCATGAACTTCCACAATTATTAAATTCAGT<br>GGTAATGCAAGAGTAAGAGGAGAGGTGGGGGGATTAACGCATTTCAGACAG<br>CTTTATAGGAGGAAAGAAAGGATTCGCGGACTTGCTGGCTGTTCGTAGTCTA<br>TGGCTGCAAAAACTACACTCTACTCACGCATACTGGAATGGTTGGCAATATT<br>CTTTTTTAGGTTAACCGGCCGGACAGTAATATAGTAATCGTTTTGTACGTTTC<br>TCAAGAAGAGACGAACAACTGTTTTGCATTTATTTTTTTTTCTTTTCCGTGAT<br>CGTCGACCATTAAAAAAATTTTTCATTCGTCGCTTGCGTGCTTCCTGGATATA<br>TAAAACACGAGTTATTTAGTTGTCTTTGTCAATTCTCAACTTTCTTTATTCATTA<br>TTTCAATTCCGTTCTTTTCATCTCTAGAACAACACAGTTACTACAACAATCAAT<br>CAATGCAGATTTTCGTCAAGACTTTGACCGGTAAAACCATAACATTGGAAGTT<br>GAATCTTCCGATACCATCGACAACGTTAAGTCGAAAATT |
| promRPL3_variant8248101 | TTTAGCACGCGGGGTGTAACTCAACAGAAAAATGTGCCATAGAACAAGACTA<br>GGCAAAAGCCAAGGAGCGTTTGCCATGAACTTCCACAATTATTAAATTCAGT<br>GGTAATGCAAAAGCAAGGGGAAAGGTGGAGGGATAAACGCATTTCCGACGG<br>CTTTATAGGCGGAAAGAAAGCACTCACAGACTTGCGGCCTGTTCACAGTCA<br>TTCGTTGCAAAAACAAAACTCTACTCACGCACACTGGAATGAATGGCAATATT<br>CTTCTTTAGGTTTACCGGCCGGACAGTAATATATTAATCGTTTCGTACGTTTTC<br>AAAGAAGCGACGCACAACTGTTTTCCATTTTTATTTTTTTCTCTTCGGTGATC<br>ATCGTCCTTGAAAAAATTTTTTCAGCTGCCTCTTTCGTGCTTCCTGGATATATA<br>AAATACGATTTATTTAGTTGTATTTTTCAGGCCTCATTCTTCATTCCTTGTTATT<br>TCATTTCGTTTTTCCTCATCTCTAGAACAACACAGTTACTACAACAATCAATCA<br>ATGCAGATTTTCGTCAAGACTTTGACCGGTAAAACCATAACATTGGAAGTTG<br>AATCTTCCGATACCATCGACAACGTTAAGTCGAAAATT |

**Table S7.** List of DNA fragments used in constructs for the generative modeling experiment.

| construct | sequence |
|---|---|
| promoter_TPM10_05 | TTTAGCACGCGGGGTGTAACTCAACAGAAAAATGTGCCATAGAA<br>CAAGACTAGGCAAAAGCCAAGGAGCGTTTGCCATGAACTTCCA<br>CAACTGCAACGCAACCAGCGCTGACGACATCCAAAGATCAAGA<br>ACTCACCCCATTCCAGGTTTCATAAGTTGGCAATATCGACATCTC<br>AATTCCCAGCGGATCTGTACTGACTAAGCTGTCTGCAGGCTACA<br>ATCAGAATTAATTTCATAGAAAAATGATGGGGAATTTCATTATAGG<br>TTAATTGCAAACACAATTAAACTTTGCAGATATTATGTCTTATTTCT<br>TCTTAAAATGGACTCTATGATTATCGTATTTTCCCACTATCGTAAC<br>ATAGGGCACCGAAAAAATAGCGATCTGTATAGATTACGTGAGTTC<br>AATATTAGCTAAGCTGTGCACATCACTTTGCCGCTTTTGTTACCA<br>TCCGCTTCTCTAAAATTGAATTAAAAAGGTGCTTTTGAAATTAATG<br>CAGATTTTCGTCAAGACTTTGACCGGTAAAACCATAACATTGGAA<br>GTTGAATCTTCCGATACCATCGACAACGTTAAGTCGAAAATT |
| terminator_ TPM10_05 | GATCCCAACGAAAAGAGAGACCACATGGTCCTTCTTGAGTTTGT<br>AACAGCTGCTGGGATTACACATGGCATGGATGAACTATACAAATA<br>GAATCATCTCTTCAAACCACAAATATTTAACGGCACTACAAAAAA<br>AGGCGCAAAAACGAAAAAGTACTACAGCAGCCTTCAAACACATT<br>CAAAAATACGCTCATGTGGATCTCTTTTTATCATCATAATGGGCGA<br>CTATTTGTCCTTCTATTTATTTTTTTTCAGACTTCTGCATCCCTTTT<br>GCGGTATGCCATATTGTTCGTCTCCATAGACAAAGAGGGGGAGA<br>TAGAACCCATCACTATCCTGTTCATTTTCCATTTATTTTTTTTCATT<br>GATTCGGCGTTATAGTAAAGAACATCAACAGGGAGGACATGTCC<br>AATATTTTACTGGAACGGCTTTGGTAAAAACTGGGGTTAGAGCCT<br>GAAATTTTCTCTTACCTGAATCTAAATAGTAATGATATGGTTGTCT<br>GTTGTGGGGAAAATGTCAAACCATAAGTTTAAAAAAGTAAAAAAC<br>GATCTTCTCAAGAAGGAATAATATAAAGCATGATAAAGTGAGCCT<br>TGGCACACGACTAGCGCTTTCAGATATTAAAAAGTTTAGATGTAG<br>GTTTTAGCGGTAACAGTTATATAAATCGTGTTTCTTCTCTTGATGA<br>AA |
| promoter_ TPM10_06 | TTTAGCACGCGGGGTGTAACTCAACAGAAAAATGTGCCATAGAA<br>CAAGACTAGGCAAAAGCCAAGGAGCGTTTGCCATGAACTTCCA<br>CAAACTTCGTATAAGCAATGAACAATCCCAAAACATACAGTAAAG<br>CGATTCTATTATATACGGGAAAGAATATCACTGCGCATGATAAAAA<br>TTTTACTGCGATAGCAAGATTCATTAATATTTTTCAACTTCTTGATT<br>GGCCCTTCGTTGTAATACAGGTGCAATGTGCACATCATTGAGGC<br>ACAAATAGGTAAAAAGTCCAAAGTGGATTCTTCTGCTGAGTGGT<br>TGATTAATAAATACGAAGCTTTCTGAACTGATTCCTCAGGTTCCA<br>ATTTTCCCATTGCTTTTTCTTTTTTTCTAATATTTTGTTATATTATTT<br>GAGTTATGCGATCATTGTGCATCTCCTTCAATTTACTACACTATTT<br>TTTTATTAAAAACAATGACCACATTCCCACGGCACCTTACCAATA<br>CAATGCAGATTTTCGTCAAGACTTTGACCGGTAAAACCATAACAT<br>TGGAAGTTGAATCTTCCGATACCATCGACAACGTTAAGTCGAAA<br>ATT |
| terminator_ TPM10_06 | GATCCCAACGAAAAGAGAGACCACATGGTCCTTCTTGAGTTTGT<br>AACAGCTGCTGGGATTACACATGGCATGGATGAACTATACAAATA<br>GAAATTATTGAATATCGGGAAAAATTTGCATAACTGAATAATTGCA<br>TATTTTCAGCAAATAAAGAGTAGGAGCGGTAGACTCGAAAAAAAT<br>ACAACATAAGTTTTTGATATGTATCCCAACCACTTGCATTCAAGTC<br>TTTTATATGAATGTCAATATGTGTTGACTTAAAGAATCGGATAGTAT<br>CTTATTCTTGATCAAGGAATAATAATATTTTCTGTCTAAAAATTAAG<br>TTCAGAAAGCGTCCTTTTGGTCACACGACTAGCGCTTTCAGATA<br>TTAAAAAGTTTAGATGTAGGTTTTAGCGGTAACAGTTATATAAATC<br>GTGTTTCTTCTCTTGATGAAA |

| | |
|---|---|
| promoter_ TPM10_07 | TTTAGCACGCGGGGTGTAACTCAACAGAAAAATGTGCCATAGAACAAGACTAGGCAAAAGCCAAGGAGCGTTTGCCATGAACTTCCACAACTTGTTGATAATTAATTCGTTGATGCAAACCTCTAAAAATGAATTGAAAAAAATAATAATTTAAGCAGTGCTCGCCTTGAGACTTGGAAGAAAAAAAAATTTAATAGAATGAAAAAAAATTTCGTGTATGTCTCTATAGTCATTTGCGTCTCCTGCGTTTATGAACCTAGATTAATATGTAAGTTAAAGAGTATATCTAGAAAATAATTGTATCAAAAGAATCAATAGCTCTATATCACTGTTTCTAATTGTATGATTTTTATATTGAGAAAAATCACCATTTTTATTTCTGATGATAACATTGCAATGTAGAAAAGCGTTGTTTTCAAAAAGCGCAATAAAAAAATCAGAAAACCCACTATGCCATTCAAAAAATAGAAAGAATATAAACGAATTCAAATAGTCCAATGCAGATTTTCGTCAAGACTTTGACCGGTAAAACCATAACATTGGAAGTTGAATCTTCCGATACCATCGACAACGTTAAGTCGAAAATT |
| terminator_ TPM10_07 | GATCCCAACGAAAAGAGAGACCACATGGTCCTTCTTGAGTTTGTAACAGCTGCTGGGATTACACATGGCATGGATGAACTATACAAATAGAAAGGACCGATGGCAAGTCGTATTTTATAGCATTCTTTCGTGATTTTTTGTCATTCTAGTTATAATTGGGGCCGTGGGCCATTTACGTGTGTGAACCTACGTGCCAAAATTTAACCCCGAACATTAATATATATAAAAATAATAAAACATAATGGTCTTGGTAATTTTCCATATTTAAGTAATTTATTTTTACCTTTGTTTAATACTGACTTTGAAACGAAAAAATTAAATTTAAGATAGATAAGAATTAACACACGACTAGCGCTTTCAGATATTAAAAAGTTTAGATGTAGGTTTTAGCGGTAACAGTTATATAAATCGTGTTTCTTCTCTTGATGAAA |
| promoter_ TPM10_10 | TTTAGCACGCGGGGTGTAACTCAACAGAAAAATGTGCCATAGAACAAGACTAGGCAAAAGCCAAGGAGCGTTTGCCATGAACTTCCACAACTTATCTACAAATTCCGCAATCCCTCCAACCAATAAATTTTTTTCCAAAATTAACTGATAACGGTAGTACTATAAATAAGTTAAATATCAATAAAAAAATTTTGAATGTCCTAGAAGTTTTCTATTTTTGCTAATTATAAAAATCCTGATATAGATTTTTGTGGAAAGGAATTCATGCGTGTTTAGTTCTACATTCTGTAGTATATTCATGTATATATATTTAATCGCACCCATGCGCTCTTTCAATCACAATCCCCGATTGATTTGTATCTGTAATAAATATTTAAAAATCCTCCTACAAATAAGTGAAGTAATTAAGCAACAAGGAAAACTCCATAAAAAAATGAAGATCATATTATATTCACCTCTAATTAAATCGTAACAAATCCTCACCATTTGTATATAATGCAGATTTTCGTCAAGACTTTGACCGGTAAAACCATAACATTGGAAGTTGAATCTTCCGATACCATCGACAACGTTAAGTCGAAAATT |
| terminator_ TPM10_10 | GATCCCAACGAAAAGAGAGACCACATGGTCCTTCTTGAGTTTGTAACAGCTGCTGGGATTACACATGGCATGGATGAACTATACAAATAGAAAATTGTACAGTTGCACAATGTTTTCTTCTATGTCGGGTTGGCATCATCGCGAACTATTGGCAGCCCAGCAAACGCTATCTCGATATGCAAATTAAGAGTTTGTGATAGCTGGATCAGATATTATACTATTTGAAATTTTCTTTACTATCATGGTTCTGTTAAGTGAATTACTATTGATTCAACATAATGCAACGTTGAGTTTGAAATGTCTTTAGTGAGAGCATAGCACTTAAATGGAGACAAAATATACACACGACTAGCGCTTTCAGATATTAAAAAGTTTAGATGTAGGTTTTAGCGGTAACAGTTATATAAATCGTGTTTCTTCTCTTGATGAAA |

| | |
|---|---|
| promoter_ TPM100_15 | TTTAGCACGCGGGGTGTAACTCAACAGAAAAATGTGCCATAGAA<br>CAAGACTAGGCAAAAGCCAAGGAGCGTTTGCCATGAACTTCCA<br>CAACTACTCAATTCGCACTTTAGCAGTGTAAGTAAGAGCGACATA<br>TACGTTGCTGTAAATGTATGTATAGGACTTAAATGGACACGACCA<br>GGCTTTTGTGCAATGAATTAGTTAAGCACGACAGATCGTACATAA<br>AGCACCCCTATTTAGCTTTTTCACCACCAGCGTCGAGTTTTATAA<br>AGGTTCCTAAGCTCAGATAACTCCTAAACCTATATCACATCCAGT<br>CATTTCTAGCTTCCTTCTGGCAAATCAAAGCCCCCCCCCAACAA<br>AAAGTTTTTAAGCATTGGATTCCGGACCCCTTCCGAGCAATCGT<br>GAAATCCGAGGCAAATAAGGTAACATTCCTGAAAGCAGGATGGT<br>TAAAAATGGAATATCGCAATCGAAGTAGAAATTGCAAGAAGTATC<br>TATAATAATGCAGATTTTCGTCAAGACTTTGACCGGTAAAACCATA<br>ACATTGGAAGTTGAATCTTCCGATACCATCGACAACGTTAAGTCG<br>AAAATT |
| terminator_ TPM100_15 | GATCCCAACGAAAAGAGAGACCACATGGTCCTTCTTGAGTTTGT<br>AACAGCTGCTGGGATTACACATGGCATGGATGAACTATACAAATA<br>GCTTAAATTTAAATGTTATATATTTTTCTGCTGTCTTTACAATTGTC<br>TTTTATCTTGTTAAGGCACCGAGGATATGATGGAGGAAGTATAAA<br>AAAAATAGAAAAGCAGTGGAGATAGGATAAAAATGTCGAGATGG<br>AGAAAATTTCAAAGCAACAAAAACTTCGATGGGAATCGAATGAA<br>GCGCCGACTTTCCTGACATCACGCCCTACATTTATCACCGTAGA<br>ATGAGGGAACCTGGGCCACAAAATTGCACACACGACTAGCGCT<br>TTCAGATATTAAAAAGTTTAGATGTAGGTTTTAGCGGTAACAGTTA<br>TATAAATCGTGTTTCTTCTCTTGATGAAA |
| promoter_ TPM100_17 | TTTAGCACGCGGGGTGTAACTCAACAGAAAAATGTGCCATAGAA<br>CAAGACTAGGCAAAAGCCAAGGAGCGTTTGCCATGAACTTCCA<br>CAACAGACAAAATTGCAGTCAAATAAGTCCTTCTCAAGTACAACG<br>CACTGAAATAATATCTTAGTAATTGAAACTATGATGTAACCACCTT<br>GTTAGAGTCTTGATATTTATAGCGGGATGTATAAAACAGCATAAAA<br>AAATGATTTTTTTTTGAAGGCTGTATTATCTTGAAATTTTTAATATT<br>TTAATGAGAGAAGGTCAAAAAAACAAAGTAGGACTTTGAGATAA<br>GTTTTCACATGTAAATATTTCCCCTATACAACACGTCATTGATCGA<br>CACCCATCTTTTTGTTCTCAAGTACGTGGTTTGCCAGCTACAACA<br>CGGTCTCAGCTTTCTTTTCCAGGCAAACAAAAAAGAAAAAATAA<br>CAAAATAAATAATTATTCATTGCTATTATACTTGTAAAGAATCATACT<br>CTCATAATTCCACACTCGCTTGTTTCTGAAAGATACCACAAACTT<br>AAGTCACTAAAAAAATGCAGATTTTCGTCAAGACTTTGACCGGTA<br>AAACCATAACATTGGAAGTTGAATCTTCCGATACCATCGACAACG<br>TTAAGTCGAAAATT |
| terminator_ TPM100_17 | GATCCCAACGAAAAGAGAGACCACATGGTCCTTCTTGAGTTTGT<br>AACAGCTGCTGGGATTACACATGGCATGGATGAACTATACAAATA<br>GACAACGTCAGCTCCATCAGCCTCTTGAACCTCACAATCATTTTT<br>TTTCTTCCGTGCAGTTCCTCAGCGCCATTTGTAGAATCTTTGTCT<br>AAAAGATAAAAAAAAATCGCAGCTGCTATAATGCAATTGAACTCT<br>AGAAAAACCTCGTATCTAAAATCATCTAGGAAATATAATCATGATT<br>GAATATATATTGCTTGAAACAGCGGATACAGTAATTAAAGGTAGC<br>GTTGGTAGCGTAGTTTTTATTACTTTGGATTCCGTGGCCACACGA<br>CTAGCGCTTTCAGATATTAAAAAGTTTAGATGTAGGTTTTAGCGG<br>TAACAGTTATATAAATCGTGTTTCTTCTCTTGATGAAA |

| | |
|---|---|
| promoter_ TPM100_18 | TTTAGCACGCGGGGTGTAACTCAACAGAAAAATGTGCCATAGAACAAGACTAGGCAAAAGCCAAGGAGCGTTTGCCATGAACTTCCACAATTCTGTCTTCTAGGGAGCATGGGTAGCCAGGCAAAATACCACACTCTAAAAAACCGCAGTGAGAAAGAGAAGGCAGGGTGGAAAGCTCAAATGAGGCCAATAACAAAAAAATTTGCTCCACGACATACCCTCGTCTTTCTCTAACTATTGAAGAAAAGGTGTAAATTTACCGGTATTGCCTTACGATATAAGAACCCGGCAAATCAACACTAATTGTAATTGGCTGTTTCCTTCAATAGTTGAGTTATCGTGCTGTCCTTACTAAGTAAAAAAAACATTATTCGTTTTTCCTCTAAAGAGCAGATGTCAGCTTGTTGCAGTGATTCTCAATTGGCGATAGATTTCTTACTTGCTTTTATTAACTAAGAATTATCGATCCAACCTCTAAACAATAAGGAAAAAAAATGCAGATTTTCGTCAAGACTTTGACCGGTAAAACCATAACATTGGAAGTTGAATCTTCCGATACCATCGACAACGTTAAGTCGAAAATT |
| terminator_ TPM100_18 | GATCCCAACGAAAAGAGAGACCACATGGTCCTTCTTGAGTTTGTAACAGCTGCTGGGATTACACATGGCATGGATGAACTATACAAATAGTATATGTGGTAAGTTGAGCCGAATTCCTTCTATGTTTGTGCAGTTTTGACGCTTATGTAAGGAGTGTAAACATTCTAGTTGTAGGTCTACGGGGCGCTAGTCTGTATCTCTCTGGCTTAGTATCGTTAATTTTCTATGATCTTCGTTACTCCTTCAGTGGTTATACGTCATATGTATTCAAGAAGAGTGTTAGATAGCTATATGCGTCGAATTAACTTATTATCTATATTTTCTTTTTTGTCGATCTTCTTGATTTAGCATATTTTAAATTTGAAAGTGGTCTACAGTGAATAAAGAGAAGGCAAATATAAAAGAAGATGCTTGGCTTTTTTTTTGTCGGCTTTTGTATCTACTAATTACGCATCTTAGTTCACATTATCATTTGGTAGAACTATACGAGATGAATCCAACAGCTTCTTCGAGGTTCAGATAATTAATAAGCTTTGATACCACATGAGAATCGTCCAATTTGTCTCATAATGGGAGCATTCGACAATGCCCACACGACTAGCGCTTTCAGATATTAAAAAGTTTAGATGTAGGTTTTAGCGGTAACAGTTATATAAATCGTGTTTCTTCTCTTGATGAAA |
| promoter_ TPM100_19 | TTTAGCACGCGGGGTGTAACTCAACAGAAAAATGTGCCATAGAACAAGACTAGGCAAAAGCCAAGGAGCGTTTGCCATGAACTTCCACAAAACTAGCTAAGCAAGTAAAATCAATCCATGAATTTAATATTAATTTTTCCTTTCTAAGTAATGACTAAATTTTCTTTTCTTTGATCTTTTCCATTTAATGATTTTTTCTTAGTTAGTTTTTCTCGTTTTACTAACTTTTATTTCTCCATCTTTTCACCCATATTTTTTATAGGCGAAATCACAAGTGTGTTCCACTTTTTCTCGTGATGAAGTTTGTTGAACTACTTTTCTGGCTGATGACCTCGACACAACCAATCATCCCCCAAAGAGCTCTTCCAAGAGTCCCGTAATTTACATGTTTTTTGTGTTAATTTTTTTAAGCAAGCAATTATATACCCCTTCACCATATTGTCTTTTCTCTACAACAGTTTCAGATATCAACAAAGTGGAAGTGTCAAAAAAATGCAGATTTTCGTCAAGACTTTGACCGGTAAAACCATAACATTGGAAGTTGAATCTTCCGATACCATCGACAACGTTAAGTCGAAAATT |
| terminator_ TPM100_19 | GATCCCAACGAAAAGAGAGACCACATGGTCCTTCTTGAGTTTGTAACAGCTGCTGGGATTACACATGGCATGGATGAACTATACAAATAGTATTTACACTACGAGAAAGACTGAAAAAAAGAAATAATTAATTCTGCTTTTAATTTTACATTTGTTAAATATTTATTTTTTCGTTTATTTATATGTTTAAGGTTAGTATATTTCGAACAATATTTGAAACATAAGTAATCCTACTCATACGCCCCTCTTTACATATATGAACTGTAGAAGGAGACGTGCATATTTGTACATAGTCATGGGACTAAATGGTAGCCAAATAACGAACGAGTAGGAATTGCACCCATTTCCAGGTTGAATGGGGGTGGTATTTGAGTCAGCCGTCCATATTTCGTGCTTGCTTGACTTGTTAAAGTAATTGAAACAACTAGAAAGAAATAGAGCAAAAGTTACAATATTTTGTTGATCAATTTCCCCATGATGGAAACATGTTAGTGAGCTATTGACATTGTTTTATACTAACTGTTAACTACTAATTATCTATATCCAATCATATAGATATCAATATACAAAGTTTGTATAAGATTAAAAGTCACACGACTAGCGCTTTCAGATATTAAAAAGTTTAGATGTAGGTTTTAGCGGTAACAGTTATATAAATCGTGTTTCTTCTCTTGATGAAA |

| | |
|---|---|
| promoter_ TPM100_20 | TTTAGCACGCGGGGTGTAACTCAACAGAAAAATGTGCCATAGAA<br>CAAGACTAGGCAAAAGCCAAGGAGCGTTTGCCATGAACTTCCA<br>CAAATCCAAAAGAAGAAAGAAAATCGAGTGAGAGCGCCTTCTAT<br>AGAGTGTAACTGCACAGACTCCGCGGAACTAAAAGAGGCATTG<br>GAAAAATGGTAAAAAGAGAAATATTGCAACAGCGTCCATTAGACC<br>CTCTGGACGAATTTTGACTCAAGAGAGCATAAAAATTACATCGAA<br>TACGTAAAGAAAGTTCTGTTGGTTTTTTTATAATCCTGTACTTTTT<br>TCATGAAACTCTCGCAGCATTGCAAAAGTAATCTTGGGATTTCTT<br>TTGCTTCGATCTCAATGGAATTAGTTATGGTCTTGCTATAGCCGC<br>GGGACAAAAAAAAAGTAAATATCGATGAAAGGGTTGTTAAAGTTA<br>AAAGGAGATTAAGTAATATTTGTGATGAAAGTTACTTATCGAGCAA<br>ATGCAGATTTTCGTCAAGACTTTGACCGGTAAAACCATAACATTG<br>GAAGTTGAATCTTCCGATACCATCGACAACGTTAAGTCGAAAATT |
| terminator_ TPM100_20 | GATCCCAACGAAAAGAGAGACCACATGGTCCTTCTTGAGTTTGT<br>AACAGCTGCTGGGATTACACATGGCATGGATGAACTATACAAATA<br>GCGGCATTGAAGAAGTACATCTGACGAAAATAAAAAAAAATTGTG<br>TATATATATTTAGGTGCCGTTAACCAGCAGCACGCTGATGGCAAG<br>TTTAACTGGGGAAAAGATGTTAGGAACATTTTGTCTGGTAGACAA<br>TGAAAAAAATAATACACATATTTATAGTCACACCCTACAATAATTGG<br>TCATAATCATATTATTTTATTCATAATATAACGCCAGATCATAATTGT<br>TTTTGTGAGAATAATTTGAAAGGGGTACATCCATCTAGAACACAC<br>GACTAGCGCTTTCAGATATTAAAAAGTTTAGATGTAGGTTTTAGC<br>GGTAACAGTTATATAAATCGTGTTTCTTCTCTTGATGAAA |
| promoter_ TPM100_23 | TTTAGCACGCGGGGTGTAACTCAACAGAAAAATGTGCCATAGAA<br>CAAGACTAGGCAAAAGCCAAGGAGCGTTTGCCATGAACTTCCA<br>CAACAGCTACTGTATATACTGAAAACATTTGTTTTAATTACCAAAA<br>AGGTCTTATTTTGAACAGATAATGAGCTTTCCATTTGCAAATGCT<br>CAATATTTGTGATCCTGTTGATATCTTCCATCTTGAAACCTTGTAT<br>AGTGATAGTATAGATCTTGCTCACATTCAATTTGAAAAATACATAC<br>ACATTTTCTCTAATTCAATACAAGTTGCTGGAATCTTCGCAAATGT<br>ATGAACAATCAAGGTGACACAGTATTTCCTGATATAACAAAATAGT<br>TTTCTAAGATTATTTTTTATCCCATTCGGCAACCTTAAGAGGTGTT<br>CCAAACGAATGCAGGTGACGGTTGTCTGACCACAAGGGAGAGA<br>TAAAGACAATAAAAAGGATCACCCAATAAGAACTTTAGATCATAAA<br>ATTTTTTTAAATCTAAAAAAAAACACATACTTAAAATTCGTTTAATA<br>AAGATATCATAGCATTAAAAAATTCTTCCTAATCAACCTGTGAAAA<br>ATGCAGATTTTCGTCAAGACTTTGACCGGTAAAACCATAACATTG<br>GAAGTTGAATCTTCCGATACCATCGACAACGTTAAGTCGAAAATT |
| terminator_ TPM100_23 | GATCCCAACGAAAAGAGAGACCACATGGTCCTTCTTGAGTTTGT<br>AACAGCTGCTGGGATTACACATGGCATGGATGAACTATACAAATA<br>GATTCTTTACATTTTTTGATGTCCTTCATATATATATGGACATTGTT<br>TAATTTTTTTTACTTTTCTTATATAGATTTTATTGTATGCGTGGATTA<br>CCGGGATGGAGTTGTCAGTGGTAATATCCATAGTAGTGGAGGTA<br>AAGGAAAGGATGTAACATATTAAAACATTTGCGCAATTAAGAACAT<br>CTTTCTATTGGGCACACTTGTATACTATATTTATATATACAAAAAAA<br>AACATTGACTTGATATCAGGTTATTACTAATTTGATATTATCGAATT<br>TTTACTACTAAAATAATCCACTAAGGTGAATTTTAATAAAGGTGGG<br>GTCTATCTATGAACACATGAAAACAAGATAGCTCAACATAGTATC<br>GAAAATAACTTAGGCTGGCATTAAGTGCTATAATAAAAGAAAGCA<br>GACCTTTGAAACCGGGGAACGACGAATGTGAAGAAGAAACATA<br>AAGCACATCACATTATCCTCTCAAGCGTTTATTGAAAGGGAGCCA<br>CACGACTAGCGCTTTCAGATATTAAAAAGTTTAGATGTAGGTTTTA<br>GCGGTAACAGTTATATAAATCGTGTTTCTTCTCTTGATGAAA |

| | |
|---|---|
| promoter_ TPM1000_26 | TTTAGCACGCGGGGTGTAACTCAACAGAAAAATGTGCCATAGAA<br>CAAGACTAGGCAAAAGCCAAGGAGCGTTTGCCATGAACTTCCA<br>CAATTGCAGCCGGTGTAGAGTTATTTTCCATACTAGATCCGTCAC<br>CCTGTTTTTACGCGCAAGAAGCAAACCAAGTTAAATACTTCACAA<br>GTAACCGCCACATAAAGTACAGAGCTAAACGAACCCTCACTATT<br>GGATTCGTCGGCCCTGCCCTATCGATGAGCCATCAGCCGACGG<br>CTTTTCAGGACTGATCGTACATTAACGGCAACCACGTCCTCCAA<br>AACTGTGTTTCTTTCTATTGACGATGTTGTGGTGTATCGGCAAGG<br>TTCTTCTTTCCACCATGGCCAAAGATCGCATCAGGAAAGAGCGC<br>ACTGGCCGAGAGGCAAGACCCCGTTCTTACCAAACTTCCTCATA<br>TAGATATGTAGCGGATTTAGGTATTAAACAAGTTATATCTTTCTTTC<br>TTACCAAACAAATGCAGATTTTCGTCAAGACTTTGACCGGTAAAA<br>CCATAACATTGGAAGTTGAATCTTCCGATACCATCGACAACGTTA<br>AGTCGAAAATT |
| terminator_ TPM1000_26 | GATCCCAACGAAAAGAGAGACCACATGGTCCTTCTTGAGTTTGT<br>AACAGCTGCTGGGATTACACATGGCATGGATGAACTATACAAATA<br>GGTTTATTTTTTATCCAGTTTAATTTGTGTTGTTATAAAATATATATG<br>TTTATTAATATTTAGTATATTTTATGTATTTTTATTTTCATTATTGCCT<br>TTTCCTTTTTTTTCGTTATGTCCTCGGGTTGTTTTTTGCTCCTGTA<br>GTCCCGCGGTGTGGTGGTTGTGACACTATTAACGCACTGGCAG<br>AAAAAATTCAAAGACTAGCCTGGTATTTTTTCGAATATTCATTTAA<br>ATCGTTCACATCTACAAACTTTCCGACTCTAGTTTTATCTGACAAC<br>ATCGTACTTAGCAAAGTTTTGGTGACCAGCAAATGTAGCAGAAC<br>AATAATTGAGCCACACGACTAGCGCTTTCAGATATTAAAAAGTTT<br>AGATGTAGGTTTTAGCGGTAACAGTTATATAAATCGTGTTTCTTCT<br>CTTGATGAAA |
| promoter_ TPM1000_27 | TTTAGCACGCGGGGTGTAACTCAACAGAAAAATGTGCCATAGAA<br>CAAGACTAGGCAAAAGCCAAGGAGCGTTTGCCATGAACTTCCA<br>CAAACCTTTTTGAGACGCCTATTCACCCACTGCAATCGCACACT<br>AAGCTGTCCAGCCTTTAACTCCGTAAGTCCTCTTCTATTTCCCAC<br>CATCCTATTATCCCGTACGCCCGGTCGGGAATCCAATGATATCTT<br>CGTAGGCGCACGCGCAAATCAGTACTTCATTTTTCCTTTTCAGT<br>GTAGCATCCATTTTCCTCTTAATGCCTTCCAATCCCCCCATCGAT<br>GATCGCAAATGAGCTCCTTCAACCTGGCAATGCTTGGGCGAAAT<br>CCGCAAACCCTCCTTTCTACCCTATTGCGAAGTCTTCCGCCTTG<br>TGAGCAATCACTAGATACGATTTTCACGATTTATTGCTTTTGTATG<br>AAACTCTTTTTAGTTCTTTTCTGTTAGTATTTTTATTCCTTAGTAGA<br>AACAAACAACCAATGCAGATTTTCGTCAAGACTTTGACCGGTAA<br>AACCATAACATTGGAAGTTGAATCTTCCGATACCATCGACAACGT<br>TAAGTCGAAAATT |
| terminator_ TPM1000_27 | GATCCCAACGAAAAGAGAGACCACATGGTCCTTCTTGAGTTTGT<br>AACAGCTGCTGGGATTACACATGGCATGGATGAACTATACAAATA<br>GTTAGTTTTTATATAGAAAAACTATTGAAAACTATAAAGAGCAGAA<br>AAGTCGTAGAATTAAGATGATTAATAATTTCTTGTTTGTTTTTGTAC<br>TTTTTTTACTTTTGCCAATTTTTGGGTAGGGACAAAATTTGTTGCA<br>GGGAGAACAGCATTCTGGTGAATTGTGTCCACCTAAGTCTTCGA<br>CGATGAGCGAAGAGGAGAATGGCCACCCATGTAGAAACTATTTC<br>ACGCATAATAATTCAAACATAGATAACAACTATTTATGTCCGTGAG<br>CGTGGACTGCAGAATTTTTATAAGCAATCACACGACTAGCGCTTT<br>CAGATATTAAAAAGTTTAGATGTAGGTTTTAGCGGTAACAGTTATA<br>TAAATCGTGTTTCTTCTCTTGATGAAA |

| | |
|---|---|
| promoter_ TPM1000_28 | TTTAGCACGCGGGGTGTAACTCAACAGAAAAATGTGCCATAGAA<br>CAAGACTAGGCAAAAGCCAAGGAGCGTTTGCCATGAACTTCCA<br>CAACAACGCGGAGGCTATTGCCTCTGCCGCTCAACCCTTCCTG<br>GCTTCCTTTTCCATTTCACAGGCGCTGTCTGAATAGGTAGGTGT<br>GAATGTTTCACCCAAAATGTGCAATGCGCGTTGGCAGATTGTCT<br>GGTTACCTTTTCACTGGTTGTTTTCCTCTTGCATCCAAATTCACA<br>TACTACTCGTTCATAGTCATTCAGCACTACCAACCAATTTTTTTCC<br>TTTCCATTCTCCGTTCGTCACTCCTCTTAAGCGTGTGCTGCTTC<br>CGCAGGCCCTCCGGCTGGGACGAGTGGGGATGATGGCCCGCC<br>CTCCTCATTGCATAAAGTCGGAAATATAATATTCCTTTGCAGGTTT<br>CCTTATCTTTTTTGATGCGTGCGTTATTCTTCATTTTTTCACACGA<br>AACATTTGTATAGATCACAACAACACAAAAATGCAGATTTTCGTC<br>AAGACTTTGACCGGTAAAACCATAACATTGGAAGTTGAATCTTCC<br>GATACCATCGACAACGTTAAGTCGAAAATT |
| terminator_ TPM1000_28 | GATCCCAACGAAAAGAGAGACCACATGGTCCTTCTTGAGTTTGT<br>AACAGCTGCTGGGATTACACATGGCATGGATGAACTATACAAATA<br>GATATCTGGAATCCACTCCGCATCATATTTATAAACTTAAAGTTATT<br>TCGAGACAGTTTCTAATTGATTACTAGTTACTTATAGGTAAAGGGT<br>TACCTTCTCATATATTTCCTTTTTAATTATGCTTATAGTCTTATATAC<br>TATTATTTTCTTTTCGATTATAAGGTGTTTAATATATGTAAAGCAAAT<br>ATATTACGGTATAAAATAATAGTCTTTTTATAGTACATGTTTTTTTTC<br>GCAATGTTTTTATTCTTCTTTGCTTTTTTCTTTCCGCCTGAGTGCT<br>TAGGTGTTAGTTCAACGGATTTTGCATTACAGTTTTTAAAGAATCT<br>ATTGCCAATAGGGGGCATATGGCTAGATATGAATGTCGCAACATG<br>TCATACTATTAGAACTTTTACTTTTCGAGCTGTTTGCTCTTTCTTC<br>TTATATGTAATCCACTCCAATGGTATCACATTATTAGGCAGCATGC<br>AAAATGAAAAAAAAACACACGACTAGCGCTTTCAGATATTAAAAA<br>GTTTAGATGTAGGTTTTAGCGGTAACAGTTATATAAATCGTGTTTC<br>TTCTCTTGATGAAA |
| promoter_ TPM1000_29 | TTTAGCACGCGGGGTGTAACTCAACAGAAAAATGTGCCATAGAA<br>CAAGACTAGGCAAAAGCCAAGGAGCGTTTGCCATGAACTTCCA<br>CAACCAAGGTTGCGCCTGCCCTCGCTCATCCCAATTTCGAGAA<br>CTTATCAAATTGAGCACATTTCACTATGGATACTGTCAGAGTCAA<br>CAATACACCCCTTTCAAACCGAACTGAAGACTTCAAATCAACTAT<br>GGAGAAAACATACTGTCAGTCACAGGCACAGCGATACTATCCAA<br>CATCCTATTATACGACATCTTATTTTTCATATTTCCATCCCTTCTTAT<br>CATTACTCCCTTCCCTTCCGTCCTCCTTTTTCCCTGCTGATTCCT<br>CCCGGGGGCCACCCCCCGGCGGGCATGTTGGAGAGAGGCGG<br>TCCCTGAGAGAATGAAAGGTAAAGATACCTGCCTAATCATGATTT<br>TCCATTTTTCGTGCTTCCTTCCACCTAACAACATATTGTATGAGCT<br>GTAAATTTTTCCATAACTTGAGTTAAATTTTATTAAATTTTTTCATAT<br>TTTTTCCCCATCTCAAGTAAATAATCTCATTTAAAAATAACAACAA<br>AAAAAGAATGCAGATTTTCGTCAAGACTTTGACCGGTAAAACCAT<br>AACATTGGAAGTTGAATCTTCCGATACCATCGACAACGTTAAGTC<br>GAAAATT |
| terminator_ TPM1000_29 | GATCCCAACGAAAAGAGAGACCACATGGTCCTTCTTGAGTTTGT<br>AACAGCTGCTGGGATTACACATGGCATGGATGAACTATACAAATA<br>GACCTAAATTTTCGTAGTTTTTTCTATTAAGTATGATATTTATTCAC<br>GGAATATATTCTTATAAGTCGATTAGGTGCAGGAAAAATGGCATTT<br>TTGGAGAATATCATAAAACATAAAAAAATTGTACAGAGACGGAATA<br>GTGAATTGTGGGAAAGCAGGGACAACAACGGTTTTCTTGCTCG<br>CAATGAGAATCTACTCCTGGCAACTTTCCACAACGAACTTTTCAA<br>GCCCCGCTATCACTTTGACCACCCCTGCGTAAGACCTGCTGCC<br>ACACGACTAGCGCTTTCAGATATTAAAAAGTTTAGATGTAGGTTT<br>TAGCGGTAACAGTTATATAAATCGTGTTTCTTCTCTTGATGAAA |

| | |
|---|---|
| promoter_ TPM1000_31 | TTTAGCACGCGGGGTGTAACTCAACAGAAAAATGTGCCATAGAA CAAGACTAGGCAAAAGCCAAGGAGCGTTTGCCATGAACTTCCA CAATGAATGCCATTGTGCCAACCATTTGCATTACTTACTATATACA GCAGTAAGCCTGATTTCAGGAGGGTACAATCCACAAGATATCCT AACTATCGAACAGATATTAAACTGTCCTTACATCATTAATCACCTT AAAAAGAGTGTTCCCTCTCCATATTATTAATTGTTCCCCCTGCTC CGTCTCATAAAACCATCACTTCAACAACTACAACAGCAATTAAGC AGACTTTTCGTCACCTTCCTACCCGGGAAACTTTGGGCAGATTT TATGCGGCCAGCCGTTTCTCACATTGTTGTTGTTGCGGGGACTC ACCAGCGGTTGGAAACTTGAGTTTTGAATAAAAACTATTGAAAAG TAGCTATATAAGAAATGAATAAAGCTTTTCTTAAAGTAAATTTAAAT TGTTTTATTCCCTTTTAAAAAGTATTTATATTTTCCTAGTTATCTTCT TTTATTTTTATTTTTATATTTTTCTAACGACCATAAAAACGAACAAC AAATGCAGATTTTCGTCAAGACTTTGACCGGTAAAACCATAACAT TGGAAGTTGAATCTTCCGATACCATCGACAACGTTAAGTCGAAA ATT |
| terminator_ TPM1000_31 | GATCCCAACGAAAAGAGAGACCACATGGTCCTTCTTGAGTTTGT AACAGCTGCTGGGATTACACATGGCATGGATGAACTATACAAATA GATAATCCTATTATGACCTATTTATTTCTTTGTTTTTTCTTTTTTTTG TTCTTTTGTACTATAAGTATAGTAATGCATTCTTGTTTTTTTAAATG CAGTTCAAATATAAAATTTATGTGACAGAATGGCAACATACTCATG TTATTCATTCAATGTACACACACACTTAAATCCACGAATCGCGAA AGTTATTGAATTCGCCCGATCGCCTTCTATCATCTGTGAAATTTTT TTTGCAATTTCCCAAGCCACTTTTTTCCTCACACGACTAGCGCTT TCAGATATTAAAAAGTTTAGATGTAGGTTTTAGCGGTAACAGTTAT ATAAATCGTGTTTCTTCTCTTGATGAAA |
| promoter_ TPM1000_33 | TTTAGCACGCGGGGTGTAACTCAACAGAAAAATGTGCCATAGAA CAAGACTAGGCAAAAGCCAAGGAGCGTTTGCCATGAACTTCCA CAAGGTCTACTGTAATTCATTTTAATACCTTTGCTCGTTACCCTTT ATCGATCCGCATGTACCGGCGATTCTCTCTCAATCAGTGCTAGC ACGGCACTGTTAAACATGATTGGTTCATCCCGATCCGCATTGTG CGTAGCAGCTGCCTCGGTGTGATTCCTTAACTCAATCTACTCCC ACACATCTCTTCATTTGTACTTCAAGAGACCGATGAGCAGTAAAA AGGTATCGGAGAGAGGCTACCCGAAGGTGGGAATATTGGGGGA GAAAAGGTGACAGGCGTTTGTTCTCTCCTCCGGCTATTCTAAAC TCCCACGGTTGGGTAGGTGAAGGAAGGAAAGCTAAAGAAAAAA AAGGAATTTTTGTCTAATGTAAGATTTTCTTTTCTTCTTTGATTTTT TACACCCAAAAAAAAATTTCTTATTAAAAAAGATTTGAGTTTTCCT TTTTTATATTTCAGAATTAATTCAAGTCATATATACAGCAAAAAACC AAATACAAAAATGCAGATTTTCGTCAAGACTTTGACCGGTAAAAC CATAACATTGGAAGTTGAATCTTCCGATACCATCGACAACGTTAA GTCGAAAATT |
| terminator_ TPM1000_33 | GATCCCAACGAAAAGAGAGACCACATGGTCCTTCTTGAGTTTGT AACAGCTGCTGGGATTACACATGGCATGGATGAACTATACAAATA GATAATCTATGTAAAAGATTTATATCTAAATATAATATCTTTAACACG TATACACAAGTTACCGATCACGTGGGGATTCTTGGGAATTGAAAA AAGTGTTTGCGCTTATGCAAGCCGCTTAGGTAAAAGGTCACCCA TTATATATGTATTAACTAAATTCATGGAATTAATCATGAAACCCAAA AACAATCAACTAAGAAAAAAACAAAACAGAAAAAACCGGAGTTTT GGATGCCAATTTTTTCTATTTTGATCCACACGACTAGCGCTTTCA GATATTAAAAAGTTTAGATGTAGGTTTTAGCGGTAACAGTTATATA AATCGTGTTTCTTCTCTTGATGAAA |

| | |
|---|---|
| promoter_ TPM1000_34 | TTTAGCACGCGGGGTGTAACTCAACAGAAAAATGTGCCATAGAA<br>CAAGACTAGGCAAAAGCCAAGGAGCGTTTGCCATGAACTTCCA<br>CAATGGAACGAGAAAGCATGTGGTCTTGAACGGTTCCTGATCCT<br>ATAATCTAAGAACATTAATTTACTCATTCACCTACTTCCACCAGTA<br>ATACCACAGGTACATATACCCTATCAACGACCTACATCATCGTAG<br>GGTCAATGGGAGTCTGCAACGCGGTCTACCCTCCAGGACAGAA<br>CCCCACCTGATCGAGGGGATAGTCGCTCACGCGTTACCACAGC<br>GGCTATGTACATTTGTCCAGTAAGGTTCTTGTACCTTCATCCGAC<br>ACCACTCACCCCCTGTTGCTTTCTCCTTGCTGCGTCACAATTTT<br>CCCCGCGGCGACTCAATGAGTGAATTAAATTGAAAATAAACTCAT<br>GTATTTTCTAGGTTTTCTTTATTCCTCTTTTGTATTGTTAAATTATTT<br>TTCCTTTTTTTCTTCCTCTAATCATTTATATTGAAAAGTATTTCATT<br>CAAGTGAGATTAACTCAACACAAAAACAAAACTATTATATATTACA<br>CTAAAAAATGCAGATTTTCGTCAAGACTTTGACCGGTAAAACCAT<br>AACATTGGAAGTTGAATCTTCCGATACCATCGACAACGTTAAGTC<br>GAAAATT |
| terminator_ TPM1000_34 | GATCCCAACGAAAAGAGAGACCACATGGTCCTTCTTGAGTTTGT<br>AACAGCTGCTGGGATTACACATGGCATGGATGAACTATACAAATA<br>GAGGATTATATATACTTTGTATTATAGTTAAAATAAGTAAGTCATTG<br>AACACCTTAGTTTATTACTATTCGCTGGTCCTCTACTGATTGTCAT<br>AGTGACTTTTATGTAATGGAAAGCATTTGCCATTCAACAGTATAAA<br>CGCTAATCTCTAGAAAAACAGACACTTTCATCGAGTAAAGGTAGA<br>GGCACTTATAAATACACAACCATTAAAGCCGATTATGAAACATGAT<br>AAAGGCACTACCTGATCCCTAACTACCAACTCCCTCTATTTTTTTA<br>CACCAAACACACACGACTAGCGCTTTCAGATATTAAAAAGTTTAG<br>ATGTAGGTTTTAGCGGTAACAGTTATATAAATCGTGTTTCTTCTCT<br>TGATGAAA |

Promoter_RPL3 and terminator_RPL3 have 40 bp homologous sequences flanking both ends for ligation with GFP and integration into XI-2 genomic site. Other promoters and terminators have 90 bp homologous sequences flanking both ends.

**Table S8.** Minimal Media Recipe.

| | **Minimal Media** |
|---|---|
| $KH_2PO_4$ | 14.4 g·L$^{-1}$ |
| $MgSO_4 \cdot 7H_2O$ | 0.5 g·L$^{-1}$ |
| $(NH_4)SO_4$ | 7.5 g·L$^{-1}$ |
| Trace metals stock solution* | 2 ml·L$^{-1}$ |
| Vitamin stock solution** | 1 ml·L$^{-1}$ |
| Glucose 50% | 40 mL·L$^{-1}$ |

*Trace metals solution: $FeSO_4 \bullet 7H_2O$ 3.0 g·L$^{-1}$, $ZnSO_4 \bullet 7H_2O$ 4.5 g·L$^{-1}$, $CaCl_2 \bullet 2H_2O$ 4.5 g·L$^{-1}$, $MnCl_2 \bullet 4H_2O$ 1 g·L$^{-1}$, $CoCl_2 \bullet 6H_2O$ 300 mg·L$^{-1}$, $CuSO_4 \bullet 5H_2O$ 300 mg·L$^{-1}$, $Na_2MoO_4 \bullet 2H_2O$ 400 mg·L$^{-1}$, $H_3BO_3$ 1 g·L$^{-1}$, KI 100 mg·L$^{-1}$, $Na_2EDTA \bullet 2H_20$ 19 g·L$^{-1}$

**Vitamins solution: d-Biotin 50 mg·L$^{-1}$, D-Pantothenic acid hemicalcium salt 1.0 g·L$^{-1}$, Thiamin-HCl 1.0 g·L$^{-1}$, Pyridoxin-HCl 1.0 g·L$^{-1}$, Nicotinic acid 1.0 g·L$^{-1}$, 4-aminobenzoic acid 0.2 g·L$^{-1}$, myo-Inositol 25 g·L$^{-1}$

The pH of the media was adjusted to 6.5 using KOH pellets. After sterilization, 2 mL·L$^{-1}$ of the trace element solution and 1 mL·L$^{-1}$ of the vitamin solution were added.

Li, Jing, Qiang Liang, Wenjiang Song, and Mario Andrea Marchisio. 2017. “Nucleotides Upstream of the Kozak Sequence Strongly Influence Gene Expression in the Yeast S. Cerevisiae.” *Journal of Biological Engineering* 11 (August): 25.

Lubliner, Shai, Leeat Keren, and Eran Segal. 2013. “Sequence Features of Yeast and Human Core Promoters That Are Predictive of Maximal Promoter Activity.” *Nucleic Acids Research* 41 (11): 5569–81.

Maaten, Laurens van der. 2008. “Visualizing Data Using T-SNE.” 2008. https://www.jmlr.org/papers/volume9/vandermaaten08a/vandermaaten08a.pdf?fbclid=IwAR0Bgg1eA5TFmqOZeCQXsIoL6PKrVXUFaskUKtg6yBhVXAFFvZA6yQiYx-M.

Nakagawa, So, Yoshihito Niimura, Takashi Gojobori, Hiroshi Tanaka, and Kin-Ichiro Miura. 2008. “Diversity of Preferred Nucleotide Sequences around the Translation Initiation Codon in Eukaryote Genomes.” *Nucleic Acids Research* 36 (3): 861–71.

Neymotin, Benjamin, Victoria Ettorre, and David Gresham. 2016. “Multiple Transcript Properties Related to Translation Affect mRNA Degradation Rates in Saccharomyces Cerevisiae.” *G3* 6 (11): 3475–83.

Shalem, Ophir, Eilon Sharon, Shai Lubliner, Ifat Regev, Maya Lotan-Pompan, Zohar Yakhini, and Eran Segal. 2015. “Systematic Dissection of the Sequence Determinants of Gene 3’ End Mediated Expression Control.” *PLoS Genetics* 11 (4): e1005147.

Teixeira, Miguel C., Pedro T. Monteiro, Margarida Palma, Catarina Costa, Cláudia P. Godinho, Pedro Pais, Mafalda Cavalheiro, et al. 2018. "YEASTRACT: An Upgraded Database for the Analysis of Transcription Regulatory Networks in Saccharomyces Cerevisiae." *Nucleic Acids Research* 46 (D1): D348–53.

Xi, Liqun, Yvonne Fondufe-Mittendorf, Lei Xia, Jared Flatow, Jonathan Widom, and Ji-Ping Wang. 2010. "Predicting Nucleosome Positioning Using a Duration Hidden Markov Model." *BMC Bioinformatics* 11 (June): 346.

Zhao, J., L. Hyman, and C. Moore. 1999. "Formation of mRNA 3' Ends in Eukaryotes: Mechanism, Regulation, and Interrelationships with Other Steps in mRNA Synthesis." *Microbiology and Molecular Biology Reviews: MMBR* 63 (2): 405–45.

Zrimec, Jan, Christoph S. Börlin, Filip Buric, Azam Sheikh Muhammad, Rhongzen Chen, Verena Siewers, Vilhelm Verendel, Jens Nielsen, Mats Töpel, and Aleksej Zelezniak. 2020. "Deep Learning Suggests That Gene Expression Is Encoded in All Parts of a Co-Evolving Interacting Gene Regulatory Structure." *Nature Communications* 11 (1): 6141.
